# XopQ induced stromule formation in *Nicotiana benthamiana* is causally linked to ETI signaling and depends on ADR1 and NRG1

**DOI:** 10.1101/2021.12.06.471425

**Authors:** Jennifer Prautsch, Jessica Lee Erickson, Sedef Özyürek, Rahel Gormanns, Lars Franke, Jane E. Parker, Johannes Stuttmann, Martin Hartmut Schattat

## Abstract

In *Nicotiana benthamiana*, expression of the *Xanthomonas* effector XopQ triggers ROQ1-dependent ETI responses and in parallel accumulation of plastids around the nucleus and the formation of stromules. Both processes were proposed to contribute to ETI-related hypersensitive cell death and thereby to plant immunity. Whether these reactions are directly connected to ETI signaling events has not been tested. Here we utilized transient expression experiments to determine whether XopQ-mediated plastid reactions are a result of XopQ perception by ROQ1 or a consequence of XopQ virulence activity. We find that *N. benthamiana* mutants lacking ROQ1, both RNLs (NRG1 and ADR1) or EDS1, fail to elicit XopQ-dependent host cell death and stromule formation. Mutants lacking only NRG1 lost XopQ-dependent cell death but retained some stromule induction that was abolished in the RNL double mutant. This analysis aligns XopQ-induced stromules with the ETI signaling cascade but not to host programmed cell death. Furthermore, data reveal that XopQ-triggered plastid clustering is not strictly linked to stromule formation during ETI. Our data suggest that stromule formation, in contrast to chloroplast peri-nuclear dynamics, is an integral part of the *N. benthamiana* ETI response and that both RNL sub-types play a role in this ETI response.

**One sentence summary:** Genetic analysis aligns effector triggered immunity (ETI) induced stromule formation with the ETI signaling cascade but not programmed cell death and questions stromule guided peri-nuclear plastid clustering.

## Introduction

Plastids exhibit exquisite developmental flexibility, as demonstrated by their capacity to differentiate into various plastid types with specialized functions, biochemical activities and internal structures, depending on the plant organ, developmental stage or environmental condition. Furthermore, plastids undergo extreme morphological changes, in some cases changing their shape within minutes or seconds (Gunning, 2005, Pyke, 2013, Delfosse *et al*., 2016). One highly dynamic feature of plastids is the projection of long, stroma-filled tubules formed by the two envelope membranes. These projections, also called stromules, are reliably observed when either the stroma or the envelope membranes are fluorescently labeled (Fig. 1; reviewed in Delfosse *et al*., 2016). Over the last two decades, stromules have been detected by fluorescence microscopy in an increasing number of plant species throughout the *Viridiplantae* (“green plants”; reviewed in Gray *et al*., 2001), suggesting that stromule formation emerged early during plant evolution. Examination of different plant tissues revealed that while stromule frequencies may vary, stromules are a ubiquitous feature of plastids (Köhler & Hanson, 2000, Holzinger *et al*., 2008).

**Fig. 1:**
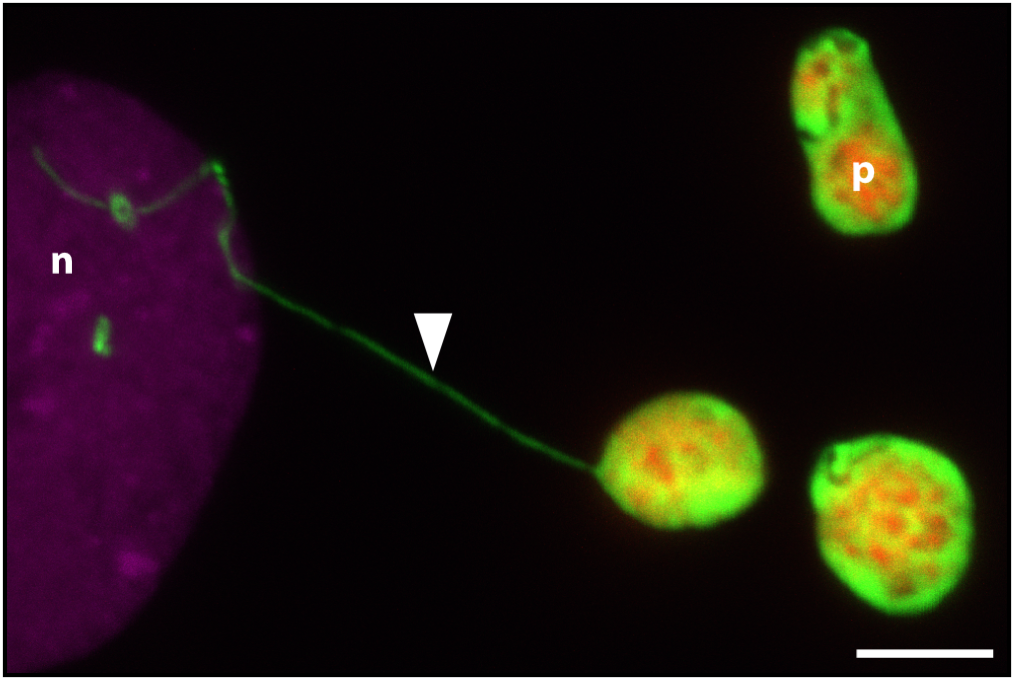
Image of a stromule connecting a plastid with a nucleus in *Nicotiana benthamiana*. CLSM fluorescence image of an *N. benthamiana* lower leaf pavement cell expressing a nuclear marker (*H2B:mCherry* in magenta) and plastid stroma marker (*FNR:EGFP* in green); chlorophyll autofluorescence is depicted in magenta; p = plastid body, n = nucleus, arrow = stromule; scale bar corresponds to 5 µm

Stromules form in response to developmental cues and increase following exposure to various stresses or signaling molecules and metabolites connected to stress (Schattat & Klösgen, 2011a, Gray *et al*., 2012, Mathur *et al*., 2012, Caplan *et al*., 2015, Vismans *et al*., 2016), suggesting stromule formation is strictly controlled by the plant. These observations led to the hypothesis that stromules participate in processes that are fundamentally important for plant survival during stress, to transmit signals and/or support physiological changes.

Despite their early emergence in the evolution of *Viridiplantae* and their frequent observation across tissues, our knowledge of stromule function is limited. To date, mutants with defects in signaling pathways regulating stromule formation were not identified. Therefore, it remains unclear which processes or functions are carried out by stromules during stress responses and how these might be executed. As an alternative to genetic dissection of stromule formation *per se*, we decided instead to test effects of mutants in defined stress responses for effects on stromule formation. Our aim is to gain insight into the role of stromules during adaptation to a specific stress and use genetic tools to decipher stromule function.

Biotic stress caused by plant interactions with recognized pathogens results in pronounced stromule formation (Krenz *et al*., 2012, Erickson *et al*., 2014, Caplan *et al*., 2015, Kumar *et al*., 2018, Adlung & Bonas, 2018). Many pathogenic microbes transfer virulence factors, known as effectors, into the host cell cytoplasm to promote infection, often by manipulating pattern-triggered immunity (PTI) programs induced by extracellular or intracellular immune receptors (Toruño *et al*., 2016, Büttner, 2016). In an incompatible interaction, intracellular immune receptors recognize one or more effectors. Effector recognition triggers a robust immune response termed effector-triggered immunity (ETI), which frequently culminates in localized host programmed cell death (a hypersensitive response = HR) at infection sites (Cui *et al*., 2015, Yuan *et al*., 2021). Most intracellular immune receptors are nucleotide binding/leucine-rich repeat (NLR) proteins. NLR proteins are represented by two major pathogen-sensing NLR receptor classes, which are defined by their N/terminal domains. So called TIR-NLRs (TNLs) possess a Toll/interleukin-1 domain (TIR) and CC-NLRs (CNLs) a coiled-coil (CC) domain at their N-terminus. Additionally, different NLR protein families have been found to function in *A. thaliana* and Solanaceous species together with pathogen-detecting (sensor) NLRs in ETI, connecting sensor NLRs with downstream immunity factors and thus are called “helper” NLRs (Cui *et al*., 2015, Wu *et al*., 2019).

Dramatic increases in stromule frequencies were observed following expression of effectors recognized by CC-NLRs or TIR-NLRs prior to ETI-induced cell death in *Nicotiana benthamiana* and *Arabidopsis thaliana* (Caplan *et al*., 2015, Erickson *et al*., 2018). For example, induction of ETI (resulting in HR) via the transient co-expression of the p50 helicase domain from tobacco mosaic virus and the cognate TIR-NLR immune receptor, N from *Nicotiana tabacum*, in *N. benthamiana* results in strong stromule induction (Caplan *et al*., 2015). Similarly, a screen by our group revealed that expression of XopQ from the bacterium *Xanthomonas campestris* pv. *vesicatoria* (*Xcv*; strain 85-10), which is recognized by the TIR-NLR immune receptor RECOGNITION OF XopQ1 (ROQ1) in *N. benthamiana*, also strongly enhanced stromule frequencies (Schultink *et al*., 2017, Erickson *et al*., 2018). In the case of ETI activation via N/p50, the authors reported that many stromules were in close proximity to the nucleus, and appeared to make contact. This observation suggested that plastids might directly deliver defense signals to the nucleus via stromules (Caplan *et al*., 2015, Kumar *et al*., 2018). Stromule frequency also increased during CC-NLR and TIR-NLR mediated ETI in *Arabidopsis thaliana* when plants were challenged with avirulent strains of the bacterial pathogen *Pseudomonas syringae* (Caplan *et al*., 2015). Thus, it appears that stromule formation is a common response of plants during ETI.

In addition to an increase in stromule frequencies and stromule-to-nucleus contacts, formation of plastid clusters around nuclei was observed during ETI responses in *N. benthamiana* (Caplan *et al*., 2015, Kumar *et al*., 2018), leading to the conclusion that plastid clusters might also support delivery of plastid-derived defense signals to the nucleus (Ding *et al*., 2019, Exposito-Rodriguez *et al*., 2020). In time lapse experiments spanning several minutes (Kumar *et al*., 2018), plastid bodies moved in the direction of stromule tips/anchor points in the majority of cases, giving the impression that stromules directionally pull the plastid body with them. This observation led to the conclusion that stromules might guide plastids to the nucleus to facilitate clustering. Hence, stromules near the nucleus might have a second function in plastid positioning.

Taken together, ETI-induced stromules present a starting point for more detailed genetic analyses of stromule formation and plastid clustering following the well-defined molecular event of effector recognition. For this, we chose *N. benthamiana* ROQ1-mediated XopQ recognition leading to ETI as a suitable system to critically examine the functional relationship between stromule formation and immunity signaling.

The TNL ROQ1 was reported to oligomerize, and assemble into a tetrameric holoenzyme with NADase activity (Ma *et al*., 2020, Martin *et al*., 2020), forming a structure similar to the pentameric complex reported for the *A. thaliana* CNL ZAR1 (‘resistosome’), which may directly integrate into membranes to function as Ca^2+^-permeable channels (Wang *et al*., 2019a, b, Bi *et al*., 2021). ETI responses initiated by TNLs like ROQ1 require at least two more components: Heterodimeric complexes composed of the lipase-like protein ENHANCED DISEASE SUSCEPTIBILITY 1 (EDS1) and either PHYTOALEXIN DEFICIENT4 (PAD4) or SENSECENCE ASSOCIATED GENE101 (SAG101), and two types of helper NLRs, which are characterized by an N-terminal four-helix bundle domain with homology to *A. thaliana* RESISTANCE to POWDERY MILDEW 8 (RPW8), and therefore are called RPW8-like coiled-coil (CC_R_) domain NLRs (RNLs) (Wagner *et al*., 2013, Jubic *et al*., 2019, Castel *et al*., 2019, Wu *et al*., 2019, Gantner *et al*., 2019, Lapin *et al*., 2019, Saile *et al*., 2020).

Molecular functions of EDS1 yet remain elusive, but its heterodimeric complexes are essential for TNL-mediated immune responses in different dicot plants including *A. thaliana* and *N. benthamiana* (Wagner *et al*., 2013, Gantner *et al*., 2019, Lapin *et al*., 2020). There are two types of RNLs known ACTIVATED DISEASE RESISTANCE GENE 1 (ADR1) and N REQUIREMENT GENE 1 (NRG1) and members of both types can be found in *A. thaliana* and *N. benthamiana* (Collier *et al*., 2011, Lapin *et al*., 2020). In *A. thaliana*, distinct EDS1-PAD4-ADR1 and EDS1-SAG101-NRG1 modules appear to regulate pathogen resistance and cell death, respectively, in TNL immunity (Lapin *et al*., 2019, Sun *et al*., 2020). In contrast, immune functions are not known for *Nb*EDS1-*Nb*PAD4 in *N. benthamiana*, and an *Nb*EDS1-*Nb*SAG101b complex appears to operate mainly through *Nb*NRG1 to mediate both cell death and resistance in this species (Qi *et al*., 2018, Gantner *et al*., 2019, Lapin *et al*., 2019). *Nb*ADR1 immune functions were so far not analyzed, but *Nb_nrg1* mutant plants, but not *Nb_eds1* mutant plants, still mobilize significant XopQ/Roq1-mediated transcriptional reprogramming, despite a loss of resistance and absence of cell death, thus suggesting some degree of cooperativity or redundancy between the two helper NLR classes also in *N. benthamiana* TNL immunity (Qi *et al*., 2018, Saile *et al*., 2020).

In this study, we capitalized on the previous characterization of XopQ/ROQ1-induced TNL immunity in *N. benthamiana* and positioned chloroplast stromule formation and perinuclear clustering in downstream signaling networks. Our data suggest that, although stromule formation is tightly linked to immune responses, it can be uncoupled from ETI-triggered cell death.

Furthermore, our data support partially redundant functions of the NRG1 and ADR1 helper NLRs (hNLRs) in stromule formation. Intriguingly, our data indicate that plastid clustering can be largely uncoupled from ROQ1 ETI and hence is unlikely to represent an integral component of the plant innate immune response.

## Results

### *Xcv*-mediated stromule formation in *N. benthamiana* depends on XopQ

We previously reported that *A. tumefaciens* mediated transient expression of XopQ from *Xcv* induces stromule formation in *N. benthamiana* (Erickson *et al*., 2018). During infection with *Xcv* strains such as *Xcv* 85-10, a strain which naturally delivers XopQ to host cells, XopQ is likely less abundant in infected cells than during transient expression experiments. Additionally, XopQ is translocated together with the entire type III-secreted effectome of *Xcv* (> 30 effectors; Teper *et al*., 2016). In order to test the extent to which stromule frequencies measured during transient expression experiments reflect the *Xcv* interaction, different bacterial strains were inoculated into *FNR:eGFP* expressing transgenic *N. benthamiana* plants. Under our greenhouse conditions the wild-type strain *Xcv* 85-10 induces an ETI-associated programmed cell death response (indicating XopQ recognition), showing first signs of dead leaf tissue at 2 dpi (Adlung *et al*., 2016). In order to be able to observe plastids in living cells, we collected leaf samples for microscopy 43 hours post-inoculation (hpi) (Fig. 2A). To test the role XopQ and other effectors play in stromule response, *Xcv* mutant strains *ΔhrcN* and *ΔxopQ* were infiltrated on the same leaf with the wild-type strain. The *ΔhrcN* mutant is deficient in typeIII secretion and serves as a non-virulent PTI control (Lorenz & Büttner, 2009). *ΔhrcN* as well as *ΔxopQ* mutant strains of *Xcv* did not induce macroscopically visual changes in infected tissues at 2 dpi and were not distinguishable from the mock infiltration (Fig. 2A; Lorenz & Büttner, 2009, Adlung *et al*., 2016, Adlung & Bonas, 2017). When analyzing the stromule phenotype, treatments differed significantly: The *Xcv* 85-10 strain induced massive stromule induction in the infiltrated tissue. The *ΔxopQ* and *ΔharcN* (no effector translation) mutant-inoculated tissue harbored almost no stromules, with levels comparable to mock inoculations (10 mM MgCl_2_) and *ΔhrcN*, which does not translocate any effectors (Fig. 2B, Supplemental materials Table S1).

**Fig. 2:**
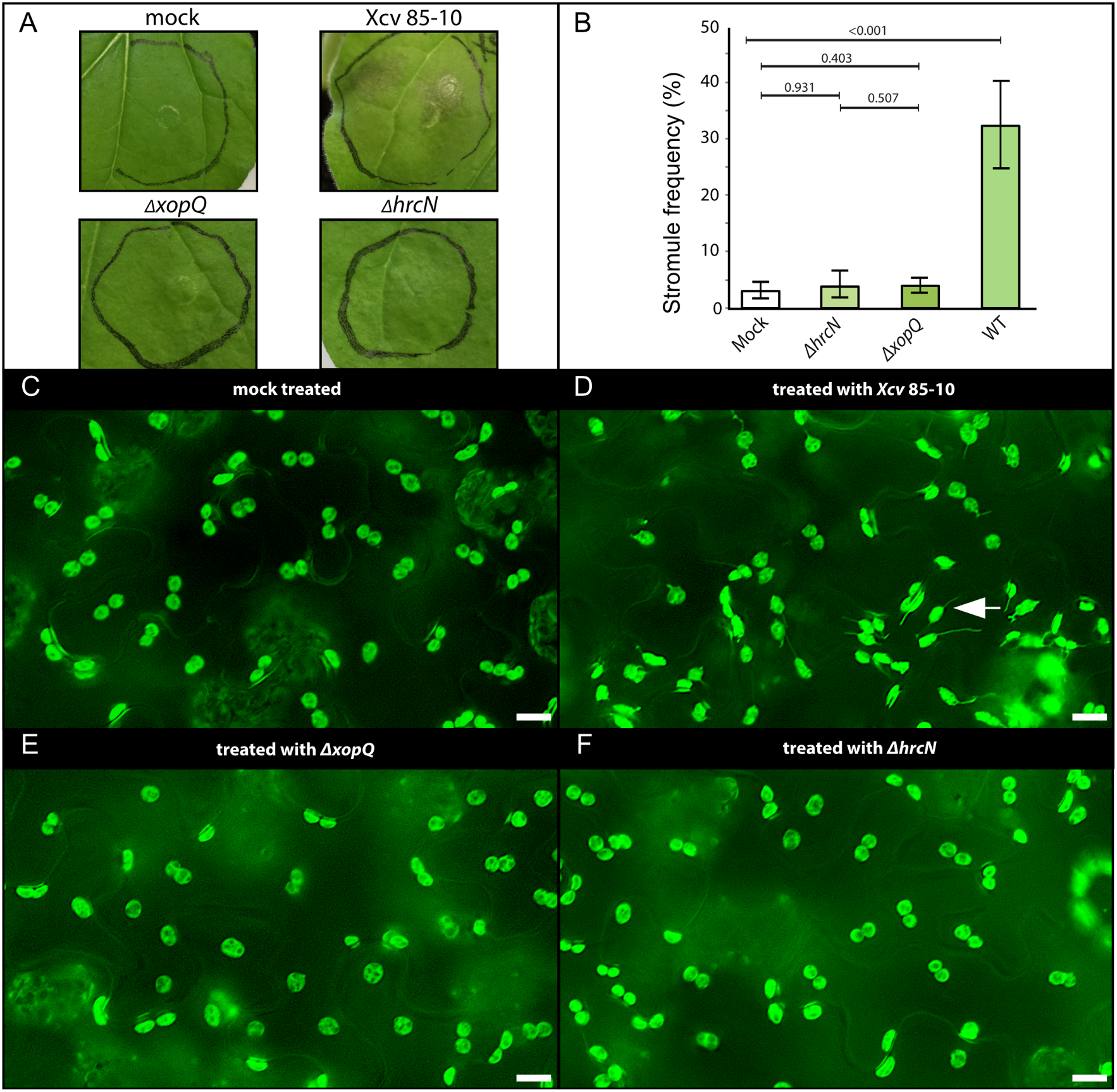
*Xanthomonas campestris pv. vesicatoria* (*Xcv*) inoculation experiments in wild-type *FNR:eGFP* transgenic plants. (**A**) Macroscopic phenotypes of leaves infiltrated with different Xcv strains at 2 dpi.(**B**) Stromule frequency (SF%) of *Xcv*-inoculated tissue at 43 hpi. Bars in the bar plot represent arithmetic averages (three repeats with 5 plants each); error bars represent 95% confidence intervals; p-values of statistical tests are shown above the black lines; (**C - F**) Sample sectors of representative microscopic images used for stromule quantification. Green fluorescence originates from the stably expressed *FNR:eGFP* plastid stroma marker; scale bars = 10 µm; arrow = stromule. (full-frame images Supplemental materials Fig. S1 and S2)

Together with transient expression experiments using *A. tumefaciens* (Erickson et al., 2018), these results indicate that stromule induction at 2 dpi by the wild-type strain is strictly dependent on presence of XopQ. No other effectors in this strain contributed measurably to stromule induction at 2 dpi, when translocated at natural levels. In conclusion, inoculations with *Xcv* strains show that XopQ-triggered stromules appear during pathogen attack at 2 dpi, and supports the idea that stromule induction during transient assays recapitulates a physiologically relevant phenotype.

### XopQ fails to induce high stromule frequency values in *roq1* mutant plants

We next examined the extent to which XopQ-mediated stromule induction in the lower epidermis of *N. benthamiana* is a consequence of XopQ perception by TNL receptor ROQ1, using *A. tumefaciens*-based transient gene expression. Different *N. benthamiana* mutant lines impaired in XopQ perception or lacking TNL downstream signaling components were co-infiltrated with Agrobacterium strains for expression of stroma-targeted eGFP (*SSU:eGFP*) and either *xopQ:mOrange2* or *mOrange2* alone, as control. Agrobacterium strains were infiltrated at a final OD_600_ = 0.2, which led to only moderate Agrobacterium-dependent stromule induction (< 20 %, Erickson *et al*., 2014) well below XopQ-induced stromule frequencies (∼ 60 %). These experimental conditions were evaluated using *N. benthamiana* stably expressing stromal-targeted eGFP (*FNR:eGFP*; Supplemental materials Fig. S3).

We first tested XopQ-mediated stromule frequencies in *roq1* mutant plants (*roq1-3* and *-4*). ROQ1-deficient plants fail to recognize XopQ and therefore lack the typical ETI-induced yellowing and chlorosis exhibited by wild-type plants following transient XopQ expression (Schultink *et al*., 2017; Gantner *et al*., 2019). *xopQ:mOrange2* or the *mOrange2* control were co-infiltrated with a stroma-targeted GFP (*ssu:eGFP*) into wild-type and mutant plants to allow for the visualization of plastids and stromules. As a control for XopQ recognition, macroscopic phenotypes of the co-infiltrated leaves were recorded 10 dpi, a time point when symptoms are clear despite the low optical densities used for infiltration resulted (Fig. 3A). As expected, *xopQ:mOrange2* expression in wild-type plants resulted in chlorosis of the infiltration spot, indicative of the XopQ-induced ETI response (Adlung *et al*., 2016). In *roq1-3* and *roq1-4* plants, there was no visible chlorosis and tissues were indistinguishable from control infiltrations. In all plant lines, *mOrange2* controls showed stromule frequency values characteristic for leaves infiltrated with "empty” GV3101 (pMP90) bacteria (compare Fig. 3B with Supplemental materials Fig. S3), indicating that the *roq1* mutation does not increase basal stromule frequencies. *xopQ:mOrange2* expression resulted in significantly higher stromule induction in wild-type plants, as previously described (Erickson et al., 2018). In contrast, *xopQ:mOrange2* expression failed to induce stromules beyond GV3101 (pMP90) basal levels in the *roq1-3* and *roq1-4* mutant plants (Fig. 3B; Supplemental materials Table S2). Average stromule frequencies in mutants infiltrated with *xopQ:mOrange2* were equal to or less than mORANGE2 controls. Fig. 3C, D and f show representative microscopy images of *xopQ:mOrange2* expressing tissues. The full loss of XopQ-induced stromule formation in *roq1* mutant plants shows that XopQ recognition by ROQ1 is required for stromule induction, and that non-recognized XopQ activity does not generate a stromule-inducing signal.

**Fig. 3:**
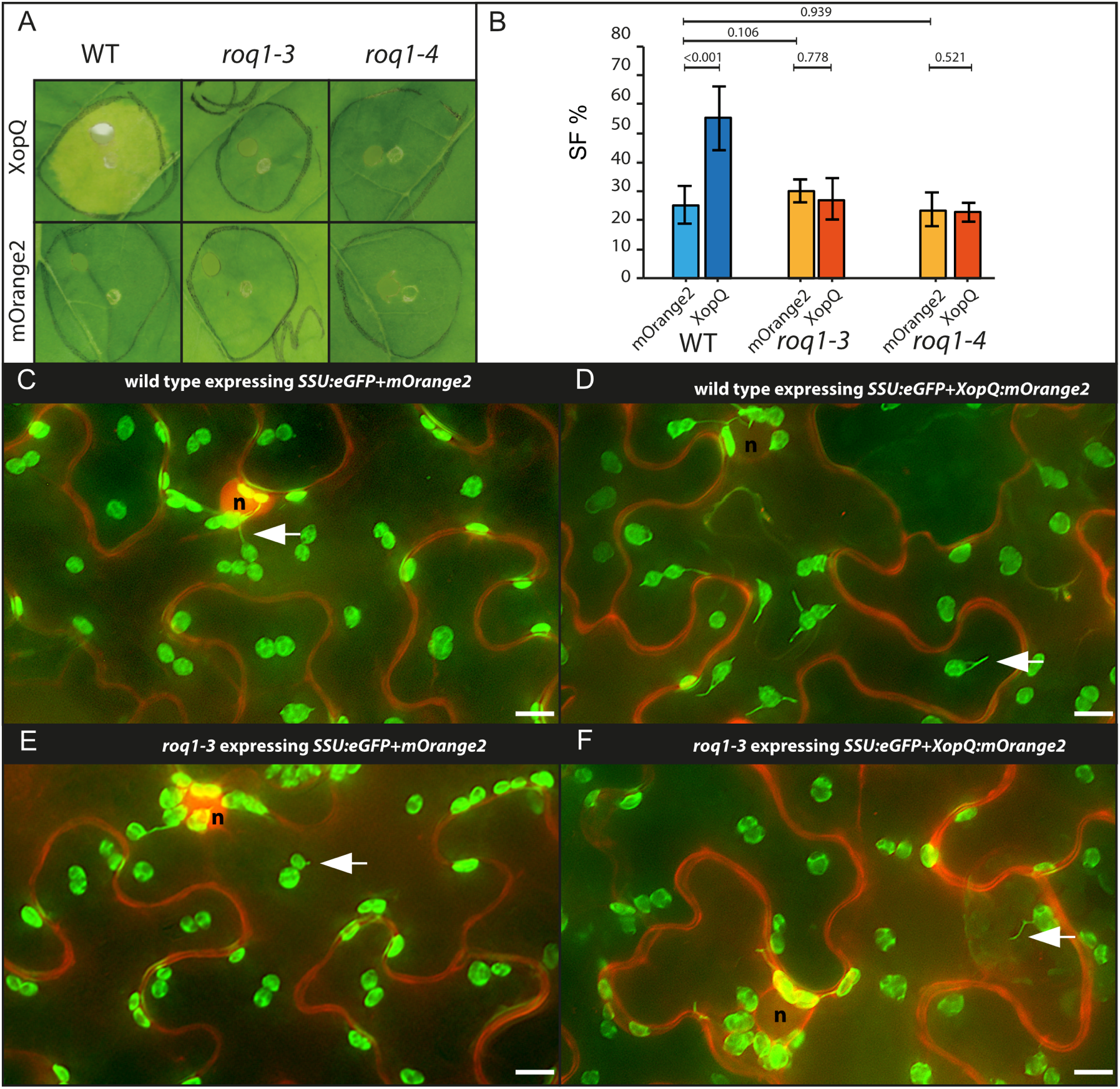
Test for XopQ mediated stromule formation in lower leaf epidermis cells of *roq1* mutants (*Nicotiana benthamiana*). (**A**) Macroscopic phenotypes of Agrobacterium-mediated *xopQ:mOrange2* and *mOrange2*-expression in wildtype, *roq1-3* and *roq1-4* leaves 10 dpi. (**B**) Results of stromule quantification expressed as stromule frequency (SF%). Bars in the bar plot represent arithmetic averages; error bars represent 95% confidence intervals; p-values of the statistical tests are shown above the black lines. (**C - F**) Sample sectors of representative microscopic images used for stromule quantification. Green fluorescence originates from the SSU:eGFP plastid stroma marker; red fluorescence originates from the mORANGE2 fluorescence protein (c and e = *mOrange2* controls; d and f *mOrange2* fused to *xopQ*); nuclei = “n”; arrow = stromule; scale bars = 10 µm. (full-frame images Supplemental materials Fig. S4 and Fig. S5)

### XopQ/ROQ1-dependent stromule induction requires EDS1

EDS1 is essential for resistance and cell death mediated by TNL-type immune receptor ROQ1 (Adlung *et al*., 2016, Gantner *et al*., 2019). In order to test if XopQ-mediated stromule formation is dependent on EDS1, mixed infiltrations were repeated in *eds1a-1* knockout and wild-type plants. In respect to stromule frequency and macroscopic phenotype the wildtype plants responded as seen in previous experiments (Fig. 4A and B; Supplemental materials Table S2). In contrast to the wildtype *eds1a-1* plants did not show signs of chlorosis at 10 dpi in response to *xopQ:mOrange2* expression, which is consistent with literature reports (Fig. 4A; Adlung *et al*., 2016, Gantner *et al*., 2019). As was the case for *roq1* mutant plants, XopQ-induced stromules were not observed in *eds1a-1* tissues (Fig. 4B and F; Supplemental materials Table S2). These results indicate that stromule formation in response to XopQ occurs downstream of EDS1 signaling, suggesting that XopQ-ROQ1 interaction and ROQ1 tetramerization (‘resistosome’ formation; Schultink *et al*., 2017, Martin *et al*., 2020) are not sufficient to induce stromules.

**Fig. 4:**
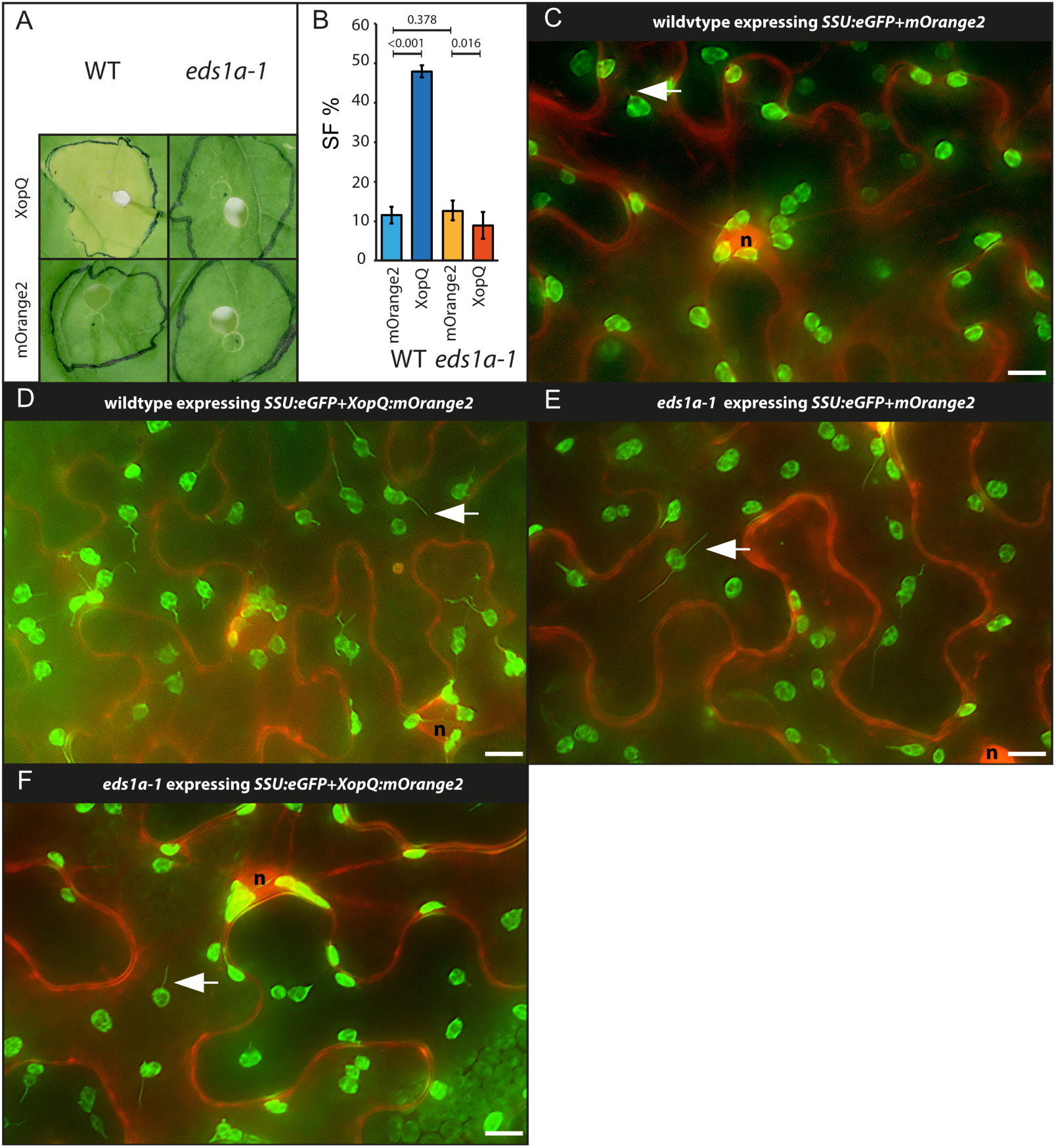
Test for XopQ-mediated stromule formation in the lower leaf epidermis of *eds1a-1* mutants (*Nicotiana benthamiana*). (**A**) Macroscopic phenotypes of *xopQ:mOrange2* and *mOrange2* infiltrated wildtype and *eds1a-1* leaves 10 dpi. (**B**) Results of stromule quantification expressed as stromule frequency (SF%). Bars in the bar plot represent arithmetic averages (three repeats with 5 plants each); error bars represent 95% confidence intervals; p-values of the statistical tests are shown above the black lines; (**C - F**) sample sectors of representative microscopic images from the data set used for stromule quantification. Green fluorescence originates from the SSU:eGFP plastid stroma marker; red fluorescence originates from the mORANGE2 fluorescence protein (c and e untagged mOrange2; d and f mOrange2 fused to XopQ); nuclei = “n”; arrow = stromule; scale bars = 10 µm. (full-frame images Supplemental materials Fig. S6.)

### XopQ-induced stromule induction depends on RNL activity but is not a consequence of host cell death

In *A. thaliana*, RNL-type NLRs of the ADR1 and NRG1 subfamilies contribute to TNL immunity (Castel *et al*., 2019, Wu *et al*., 2019, Lapin *et al*., 2019, Saile *et al*., 2020). In *N. benthamiana nrg1* mutant plants, resistance and cell death induced by several TNLs was fully abolished, suggesting NRG1 as the major RNL in TNL immunity in this species (Qi *et al*., 2018). To test if XopQ- stimulated stromule formation is NRG1-dependent, two mutant lines with different genomic deletions, *nrg1-4* and *nrg1-5* (Ordon *et al*., 2021) were analyzed. The *nrg1* mutants did not show signs of yellowing in response to *A. tumefaciens* mediated XopQ expression, and infiltration spots were macroscopically indistinguishable from control infiltrations (Fig. 5A), as expected (Qi *et al*., 2018, Ordon *et al*., 2021). All three plant lines responded similarly to the *mOrange2* control expression, with stromule frequencies reaching approximately 25 % (Fig. 5B; Supplemental materials Table S2). In contrast, the response to *xopQ*:*mOrange2* was markedly different between wild-type and *nrg1* mutant lines (Fig. 5B). *xopQ*:*mOrange2* expression in the *nrg1* background induced stromule frequencies values, which were between Agrobacterium-inoculated and x*opQ*:*mOrange2* -inoculated wild-type plants. Although *roq1, eds1* and *nrg1* mutants were equally deficient in XopQ-induced cell death (necrosis), stromule formation did not strictly require NRG1, and is thus uncoupled from cell death in *Nb_nrg1* mutant plants.

**Fig. 5:**
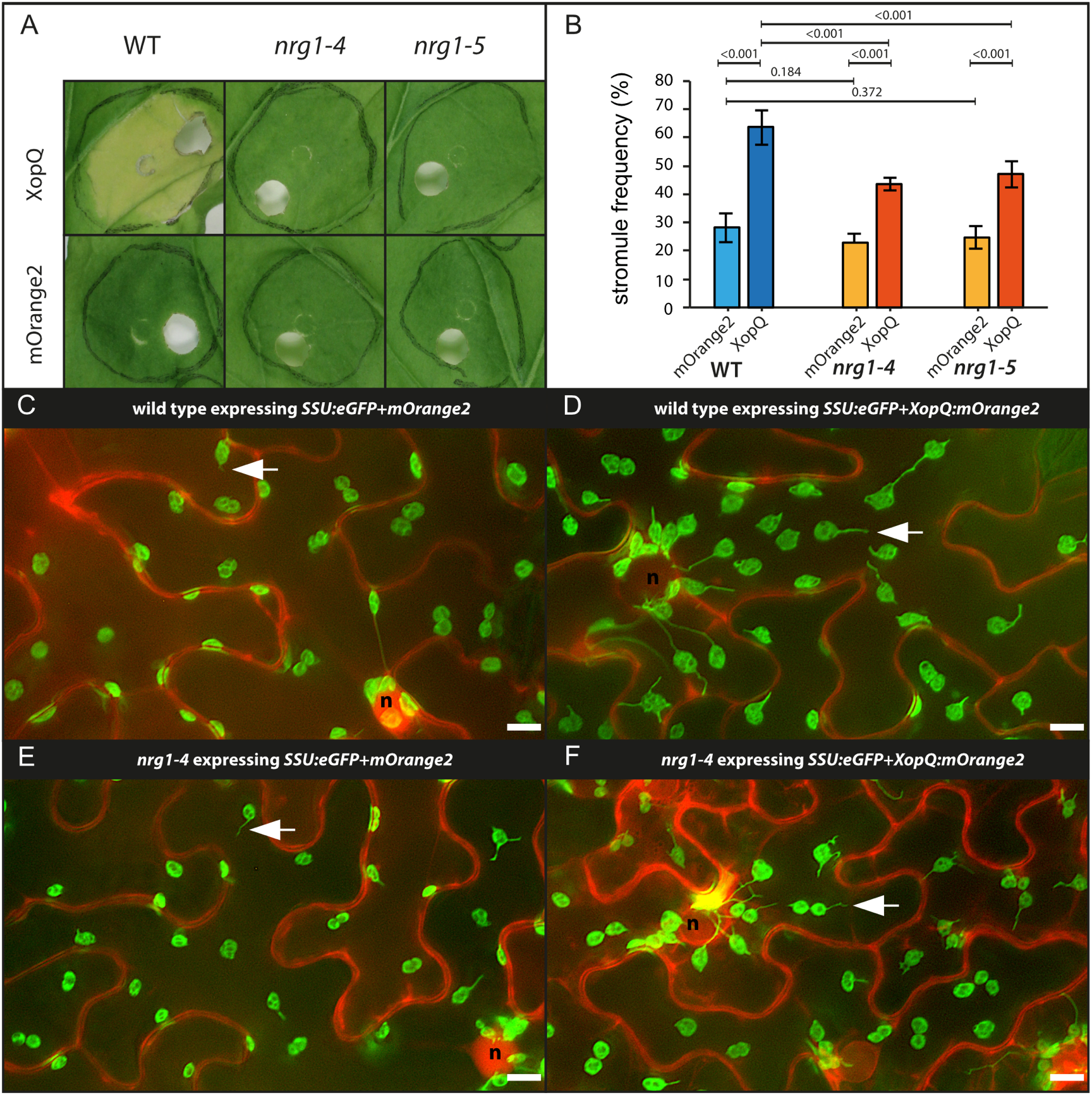
Test for XopQ mediated stromule formation in lower leaf epidermis cells of *nrg1* mutants (*Nicotiana benthamiana*). (**A**) Macroscopic phenotypes of *XopQ:mOrange2* and *mOrange2*-infiltrated wildtype, *nrg1-4* and *nrg1-5* leaves at 10 dpi.(**B**) Results of stromule quantification expressed as stromule frequency (SF%). Bars in the bar plot represent arithmetic averages (three repeats with 5 plants each); error bars represent 95% confidence intervals; p-values of the statistical tests are shown above the black lines. (**C - F**) Sample sectors of representative microscopic images used for stromule quantification. Green fluorescence originates from the SSU:eGFP plastid stroma marker; red fluorescence originates from the mORANGE2 fluorescence protein (c and e = *mOrange2* control; d and f = *mOrange2* fused to *XopQ*); nuclei = “n”; arrow = stromule; scale bars = 10 µm. (full-frame images in Supplemental materials Fig. S7).

So far, a function of ADR1 was not identified in *N. benthamiana* (Lapin et al., 2019, Sun et al., 2021) but varied contributions of these RNLs to Arabidopsis TNL ETI suggested that *N. benthamiana* ADR1 might steer residual stromule formation in *N. benthamiana nrg1* lines. Therefore, we tested stromule formation in response to XopQ in a newly generated *Nbadr1_nrg1* double mutant line using mixed infiltrations as before. As in the *nrg1* single mutants, *xopQ:mOrange2* expression did not induce yellowing or cell death in *adr1_nrg1* plants (Fig. 6A). In contrast to our observations in *nrg1* single mutants, *xopQ:mOrange2* did not induce the formation of stromules beyond the mORANGE2 control (Fig. 6B; Supplemental materials Table S2). Accordingly, the *adr1_nrg1* mutant exhibited stromule and cell death phenotypes similar to the *roq1* (Fig. 3A) and *eds1* (Fig. 4A) mutant lines. Overall, these results show that not only NRG1 but also ADR1 contributes to stromule formation in *N. benthamiana*. Hence, these results uncover that the ADR1-family of RNLs exhibits not only ETI signaling functions in *A. thaliana* ETI but also functions in the ETI response of solanaceous plants.

**Fig. 6:**
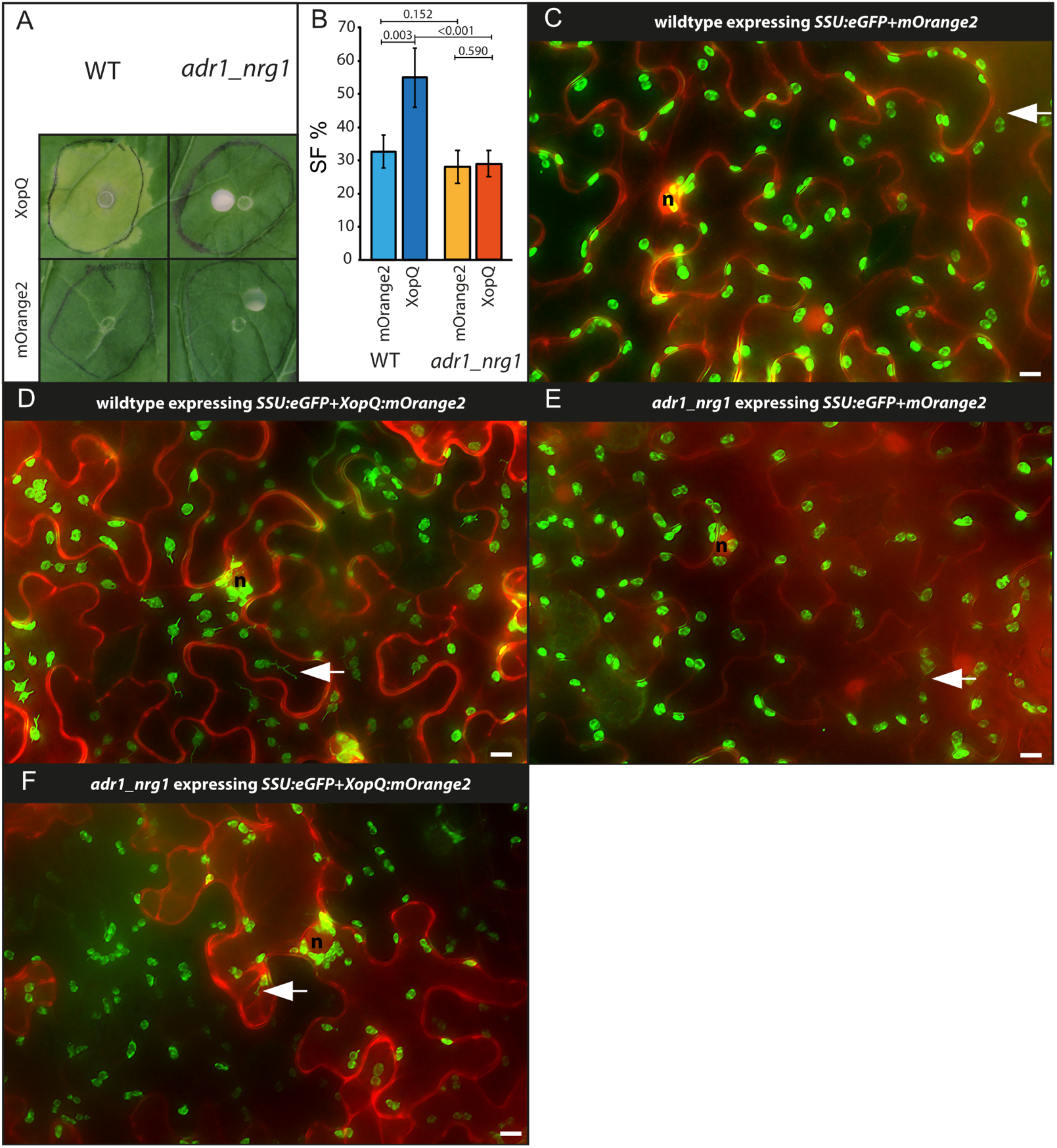
Test for XopQ mediated stromule formation in lower leaf epidermis cells of adr1_nrg1 double mutants (*Nicotiana benthamiana*). (**A**) Macroscopic phenotypes of *xopQ:mOrange2* and *mOrange2*-infiltrated wildtype and *adr1*_*nrg1-5* leaves at 10 dpi. (**B**) Results of stromule quantification expressed as stromule frequency (SF%). Bars in the bar plot represent arithmetic averages; error bars represent 95% confidence intervals; p-values of the statistical tests are shown above black lines; wild-type 3 plants and *adr1_nrg1* 5 plants for each of 3 repeats. (**C - F**) Cropped images used for stromule quantification. Green fluorescence originates from the SSU:eGFP plastid stroma marker; red fluorescence originates from the mORANGE2 fluorescence protein (c and e are *mOrange2* controls; d and f show *mOrange2* translation fusion with *XopQ*); nuclei = “n”; arrow = stromule; scale bars = 10 µm. (full-frame images shown in Supplemental materials Fig. S8).

### XopQ-induced peri-nuclear plastid clustering does not require TNL immune signaling

In order to test if XopQ-induced ETI facilitates formation of chloroplast clusters, and whether this has the same genetic dependencies as found for stromule frequency, chloroplast clustering was quantified in wild-type, *roq1-3, eds1a-1, nrg1-4* and *adr1_nrg1* lines. As a measure for plastid clustering, the number of plastids in close proximity (up to one plastid in diameter) to the nucleus was counted and expressed as the plastid nucleus association index (PNAI, see Erickson *et al*., 2014, 2018). In these experiments, mORANGE2 or XopQ:mORANGE2 fluorescence, respectively, served to highlight the position of nuclei (see Fig. 7B - K). In control infiltrations (*mOrange2*), wild-type and all four mutants produced similar numbers of plastids around the nucleus (Fig. 7A; Supplemental materials Table S3). No significant differences in PNAI were detected. When challenged with XopQ, plastid clustering increased in wild-type plants (Fig. 7A; Supplemental materials Table S3). Although plastid clustering has been considered an ETI response (Caplan *et al*., 2015, Kumar *et al*., 2018, Ding *et al*., 2019), we found that XopQ-induced plastid clustering was not diminished in mutant lines impaired in XopQ recognition or downstream signaling (Fig. 7A; Supplemental materials Table S3). Upon expression of *xopQ:mOrange2, nrg1-4* showed wild-type levels of plastid clustering, while *roq1-3, eds1* and *adr1_nrg1* mutants had significantly higher PNAI values (Fig. 7A; Supplemental materials Table S3). Fig. 7B - K shows representative images of the different plant lines expressing *ssu:eGFP*+*xopQ:mOrange2* or *ssu:eGFP*+*mOrange2*.

**Fig. 7:**
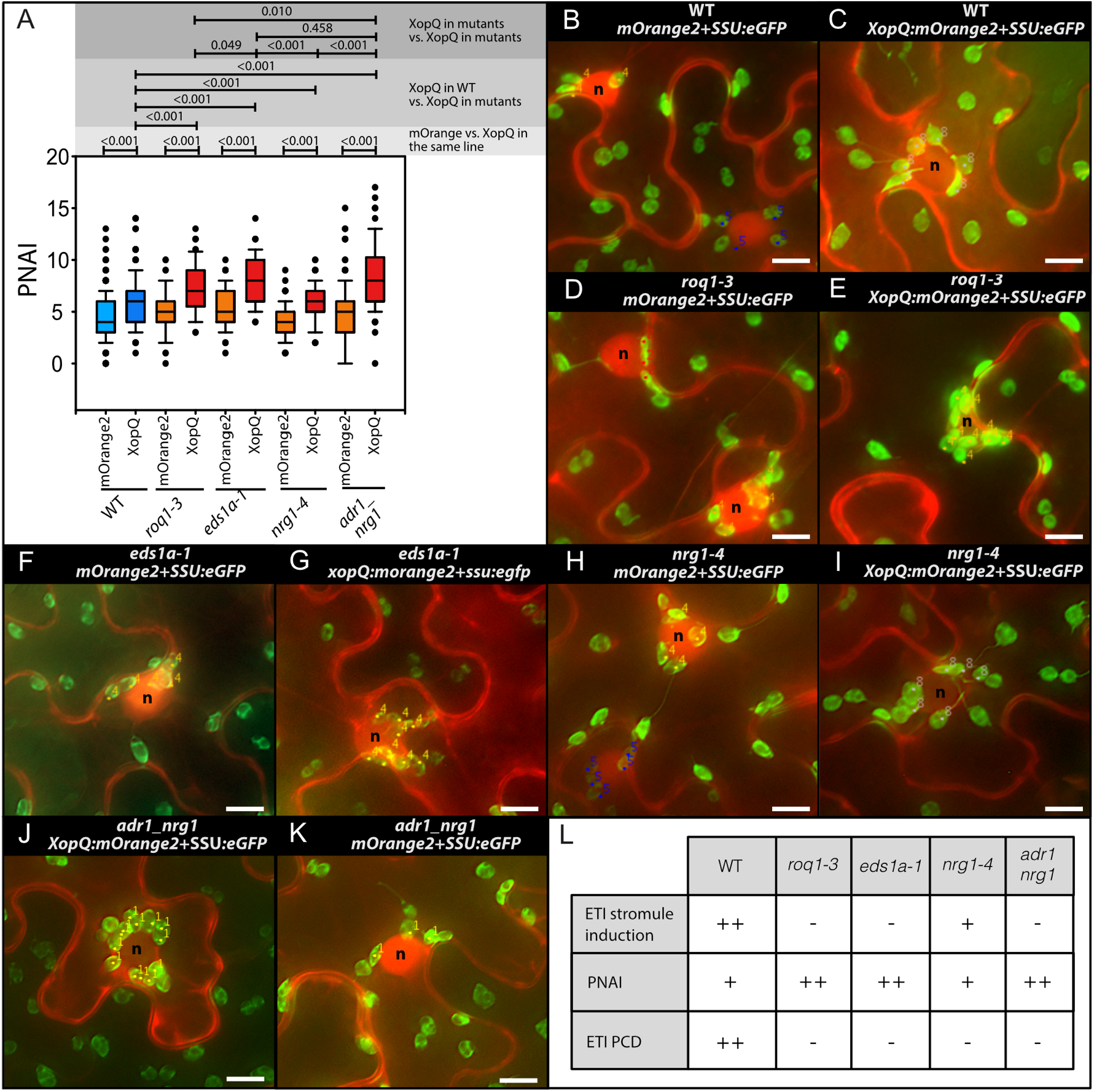
PNAI - Analysis of plastid clustering in response to XopQ expression (*Nicotiana benthamiana*). Plastid nucleus association index (PNAI) at 3 dpi following *mOrange2* and *xopQ:mOrange2* expression in wild/type as well as mutant plant lines (*roq1-3, eds1a, nrg1-4* and *adr1_nrg1*). (**A**) PNAI values represented in box plots (box line = median, whiskers = 10^th^ as well as 90^th^ percentile, with each outlier plotted); p-values of Mann-Whitney rank sum tests are shown above the lines in grey boxes. (**B - K**) Sample images of nuclei with associated plastids (labeled with a dot a number as they were during original plastid scoring); “n” = nuclei; scale bars = 10 µm. (**L**) Summary of observed ETI stromule, PNAI and cell death phenotypes where “-” = no change compared to control, “+” = visible but moderate increase in cell death; “++” = strong increase.

We concluded that when the XopQ-induced ETI signal cascade is blocked (*roq1, eds1* and *adr1_nrg1* mutants), the tendency of plastids to cluster around the nucleus remains, and is even enhanced compared to wild-type or plants (*nrg1* mutant) with partially intact signaling (Fig. 7H). These data suggest that one feature of ROQ1/XopQ-induced ETI is suppression of plastid clustering.

## Discussion

Here, we set out to understand how ETI-associated stromule formation aligns with signaling processes downstream of immune receptor activation, using recognition of the effector XopQ by the TNL ROQ1 in *N. benthamiana* as a case study. A quantitative analysis of stromule formation and peri-nuclear plastid clustering in XopQ recognition and ETI signaling mutants produced several important insights. First, complete absence of a XopQ-related stromule response in *roq1* and *eds1* mutants shows that XopQ-induced stromules are not a result of its virulence/effector activity, but result from effector recognition by the ROQ1 immune receptor and a resulting *EDS1*-dependent ETI response. Second, residual induction of stromules in the *nrg1* mutant in the absence of appreciable cell death suggests that induced stromule formation is not a consequence of ETI-related host cell death but more closely related to pathogen resistance. Third, analysis of the *nrg1_adr1* double mutant line reveals that residual XopQ-induced ETI and stromule frequency in *nrg1* (but not *eds1*), is conferred by ADR1 in *N. benthamiana*. This reveals an ADR1 contribution to TNL ETI processes in a solanaceous plant, suggesting usage of both RNLs NRG1 and ADR1 branches in ETI, as observed in Arabidopsis. Finally, ROQ1/XopQ-induced plastid clustering does not correlate with ETI induction and stromule formation and is thus likely to be a direct or indirect consequence of XopQ virulence activity during infection. A model summarizing these findings is presented in Fig. 8A and B.

**Fig. 8:**
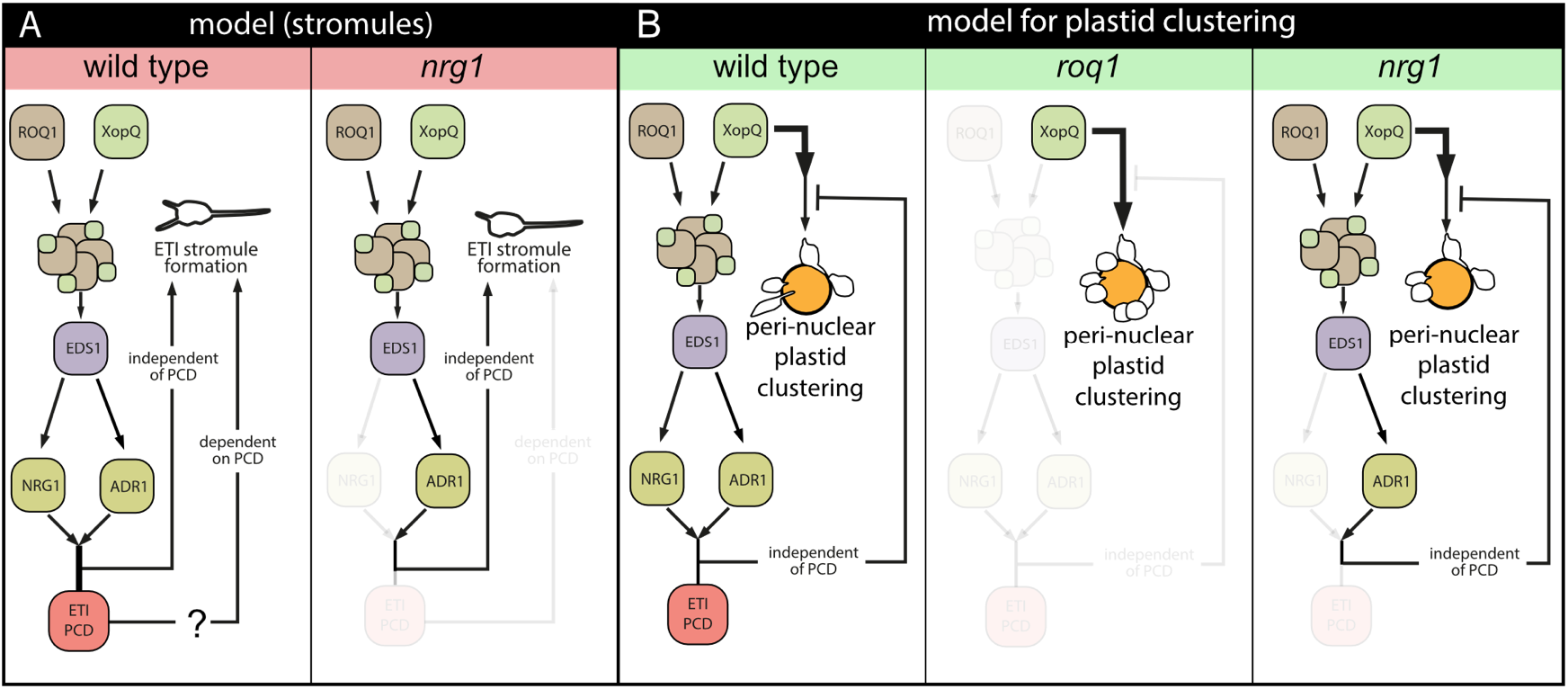
Model of XopQ mediated stromule induction and peri-nuclear plastid clustering. (**A**) Model for effector triggered immunity (ETI) induced stromule formation by XopQ in wild-type plants (left panel): XopQ (*<u>X</u>anthomonas <u>o</u>uter <u>p</u>rotein <u>Q</u>*) is recognized by the TIR-NB-LRR (TOLL INTERLEUKIN 1 RECEPTOR NUCLEOTIDE BINDING LEUCIN RICH REPEAT) resistance protein ROQ1 (RECOCNITION OF XopQ 1), which forms with XopQ a heteromeric protein complex, the so called ‘resistosome’. From the ‘resistosome’ the information of XopQ recognition is transferred with the help of EDS1 (ENHANCED DISEASE SUSCEPTIBILITY 1) to the RNL class helper NLRs NRG1 and ADR1, which culminates in PCD (<u>p</u>rogrammed <u>c</u>ell <u>d</u>eath) as well as stromule formation. In *nrg1* mutants (right panel) XopQ fails to induce cell death but at the same time still shows significant ETI stromule induction mediated by ADR1. This indicates the existence of a PCD independent ETI stromule induction pathway. However, at this point a role for PCD in stromule induction, or a role for stromules in PCD is not ruled out (?). (**B**) Model for peri-nuclear plastid clustering following *xopQ* expression: In wildtype plants (left) an intact ETI signal chain partially suppresses the strong peri-nuclear plastid clustering induced by XopQ presence. When ETI signal transduction is blocked (*eds1, roq1* and *adr_nrg1* mutants) this suppression does not take place and plastid clustering is enhanced (middle). The restricted ETI signal chain in *nrg1* mutants (right) is sufficient for suppression of induced peri-nuclear plastid clustering to wildtype levels, thus suppressing signals likely originate from this pathway. Both models only consider genes, which were analyzed as part of this study. The role of the different proteins forming distinct complexes with EDS1 will have to be elucidated in future.

### Uncoupling immune signaling and HR-induced stromules

Stromules have been proposed to transmit retrograde signals to the nucleus, and amplify programmed cell death responses as part of ETI (Caplan *et al*., 2015). More recently it was suggested that stromules have the added function of locating and pulling plastid bodies to the nucleus (Kumar *et al*., 2018). So far, all reported microbial effectors that induce stromules also provoked programmed cell death (Caplan et al., 2015, Erickson et al., 2018). Therefore, it remained unclear whether stromule formation accompanies the cascade of events contributing to host cell death or is a by-product of physiological changes in cells as they die. The observed stromule induction upon XopQ expression in *nrg1* mutant plants without cell death or measurable resistance (Qi *et al*., 2018) (Fig. 5A) suggests stromules represent events upstream of ETI-related pathogen resistance and host cell death. Hence, ETI-induced stromule formation is not coupled to cellular destruction but more likely an integral part of the ETI response, as suggested previously (Caplan *et al*., 2015).

### ADR1 contributes to TNL immunity and stromule formation in *N. benthamiana*

Dicot genomes encoding TNL receptors generally also encode RNL-type NLRs of the ADR1 and NRG1 classes (Collier *et al*., 2011, Lapin *et al*., 2020). In *A. thaliana*, both classes of RNLs contribute to immunity to different extents. Three *A. thaliana* ADR1 paralogs have functions in basal immunity to virulent pathogens related to salicylic acid, and they also contribute to PTI (Bonardi *et al*., 2011, Jubic *et al*., 2019, Pruitt *et al*., 2021, Tian *et al*., 2021). ADR1s and NRG1s contribute to different extents to *A. thaliana* TNL immunity, and function as distinct modules, respectively with EDS1-PAD4 and EDS1-SAG101 dimers regulating pathogen resistance and cell death (Lapin et al., 2019, Saile et al., 2020, Lapin et al., 2020, Sun et al., 2021). In *N. benthamiana*, only EDS1-SAG101 appeared to execute TNL ETI, and NRG1 was identified as a major RNL required for cell death and resistance mediated by several tested TNLs (Qi *et al*., 2018, Gantner *et al*., 2019). A role for *N. benthamiana* ADR1 in immunity was so far not detected, although Qi et al. (2018) reported residual transcriptional reprogramming occurring upon ROQ1 activation in *nrg1*, but not *eds1* mutant plants. Notably, *NbADR1* was among the upregulated genes of an *NRG1*-independent regulon (Qi et al, 2018). Residual stromule formation in *nrg1*, but not *eds1* or *nrg1_adr1* (compare panel ‘(A)’ in Supplemental materials Fig. S4, 5 and 6) supports an *Nb*ADR1 contribution to TNL-ETI in *N. benthamiana*. Hence, stromule formation appears to be a highly sensitive read-out for ETI induction, occurring in the absence of cell death or measurable pathogen resistance which are blocked in *N. benthamiana nrg1* and *eds1* mutants (Qi et al 2018). Current evidence suggests that, besides the sensor CNL ZAR1, RNLs also might assemble into pore-forming resistosome complexes and function as Ca^2+^ permeable cation channels (Wang *et al*., 2019a, Jacob *et al*., 2021, Bi *et al*., 2021). Induced Ca^2+^influx into host cells would then amplify ROS generation and salycilic acid signaling as well as transcription (Lu and Tsuda 2021, Yuan *et al*., 2021). In future work, it will be interesting to examine whether Ca^2+^ levels inside cells influence stromule formation, as stromule-to-nucleus connections were found to contribute to ROS formation (Caplan *et al*., 2015).

### Is perinuclear plastid clustering independent of ETI stromule induction?

In an emerging concept, plastids are the source of important immune response signaling and defense metabolites, including precursors of salicylic acid and jasmonic acid. Many plastid-derived signals must reach the nucleus to fulfill their proposed functions (reviewed in Kretschmer *et al*., 2020). Thus, re-localization of plastids towards the nucleus might promote more efficient signal transmission. Indeed, when challenged with different pathogens and H_2_O_2_, plastids relocated towards the nucleus in *N. benthamiana* epidermis leaf cells, forming peri-nuclear clusters (Erickson *et al*., 2014, Caplan *et al*., 2015, Ding *et al*., 2019a). How plant cells regulate re-localization of plastids to the nucleus upon different stimuli remains unknown. Based on observations of stromule orientation often coinciding with plastid directional movement, it was proposed that stromules are initiated during ETI and extended along the microtubules network, finding anchor points on actin filaments close to the nucleus which guide plastid body movement towards nuclei (Kumar *et al*., 2018). Peri-nuclear plastid clustering, as a consequence, might enhance plastid-to-nucleus signal transfer underpinning immune responses. When we challenged wild-type plants with *XopQ:mOrange2*, stromule formation as well as perinuclear plastid clustering were consistently induced in lower epidermis cells (Fig. 7A -K). Additionally, stromules facing the nucleus and seemingly anchored in the nuclear periphery were observed (see Fig.s 3B, 4B, 5B and 6B for SF%; e.g. 3D, 4D and 5D for nucleus-associated stromules). Both observations support stromules guiding plastid body movement (Kumar et al., 2018). However, based on this model we expected impaired or abolished perinuclear clustering in plant lines unable to recognize XopQ (*roq1*) or to initiate TNL immunity (*eds1, nrg1, nrg1_adr1*). Despite, having reduced numbers of ETI-associated stromules, the plastid clustering still occurred when *XopQ:mOrange2* was expressed in respective mutant backgrounds. Notably, plastid clustering was more pronounced in the mutants compared to wild type. In contrast, perinuclear clustering in response to *XopQ:mOrange2* expression was not slightly reduced in *nrg1* plants (Fig. 7 A-K). In summary, we observe a negative correlation between stromule frequency and perinuclear plastid clustering. Accordingly, in our assays ETI induction of stromules was associated with lower plastid clustering compared to when ETI was disabled (Fig. 7H). These data suggest that increased chloroplast guidance towards the nucleus via stromules does not facilitate ETI but might be due to XopQ virulence activity.

### Is induction of peri-nuclear plastid clustering a part of XopQ’s function?

Stronger plastid clustering observed in the absence of ROQ1, EDS1, ADR1_NRG1 suggests it may represent a consequence of undisturbed XopQ activity (Fig. 7A and 8B). This observation partially contradicts the suggestion that peri-nuclear plastid clustering supports ETI responses by facilitating more efficient transfer of pro-defense signals from plastids to the nucleus (discussed in Exposito-Rodriguez et al., 2020). If the sole function of peri-nuclear plastids is to enhance ETI, why should the bacteria facilitate peri-nuclear plastid clustering via XopQ in the absence of effector recognition? Conversely, why suppress clustering when ETI is induced by XopQ? Our results suggest that clustering may serve multiple functions or is the consequence of several stimuli in plant pathogen interactions. In support of this hypothesis while plastid clustering in *N. benthamiana* occurs in response to ETI triggering stimuli (e.g. TMV-p50 and RPS2 recognition Kumar *et al*., 2018, Ding et al., 2019), it also occurs in response to pattern triggered immunity (PTI) stimuli (*Pst* DC3000 *ΔhopQ1-1*, flg22 and H_2_O_2_, Ding et al., 2019), which demonstrates that plastid clustering is not ETI specific. Additionally, plastid clustering is not restricted to plant microbe interactions and has been found to be important for plastid inheritance during cell division (Sheahan et al., 2004 and 2020) and has been observed following the exposure of *N. benthamiana* epidermis leaf cells to cytokinin (Erickson et al., 2014). In summary, although plastid accumulation at the nucleus is linked to plant microbe interactions it is not exclusively so, and may reflect one output resulting from changes to different cell physiological parameters (i.e., altered hormone or H_2_O_2_ levels). Currently, although we see that XopQ activity induces clustering, the trigger for this phenotype remains enigmatic and it remains to be seen whether it is of any benefit to *Xcv* during an infection.

## Conclusion

We set out in this study to test two hypotheses about stromule function in the context of effector triggered immunity. We provide here experimental evidence for a direct link for ETI-induced stromule formation, supporting the hypothesis of Caplan et al. (2015), which suggests that stromules play a specific role during ETI. Our findings therefore encourage the enquiry of the nature of this specific role in future. In contrast to this, our results do not support the second hypothesis, which suggested that stromules might be needed to guide plastid movement towards the nucleus (Kumar et al., 2018), highlighting the fact that there is currently no mechanistic explanation for peri-nuclear plastid accumulation and that for this phenomenon fundamental work still has to be done. Finally, the obtained results in the *nrg1* and *adr1_nrg1* mutants (cooperative activity of NRG1 and ADR1 in ETI signaling) show the potential of stromules to function as a quantifiable and sensitive readout for branches of ETI signaling, which result in currently used assays only in hard to detect differences.

## Material and Methods

### Plant material

*Nicotiana benthamiana* plant lines used in this study were: wild type, *roq1-3* and *roq1-4* (Gantner *et al*., 2019), *eds1-1* (also referred to as *eds1a-1*; Ordon *et al*., 2017), *nrg1-4* and *nrg1-5* (Ordon *et al*., 2021). An *adr1_nrg1* double mutant was created by genome editing using a derivative of pDGE311, a plant transformation vector containing additional counter-selection markers and an intron-optimized *zCas9i* gene, as recently described (Grützner *et al*., 2021, Stuttmann *et al*., 2021) (Methods S1). For *Xanthomonas* inoculations transgenic *N. benthamiana* plants of the plant line FNR:eGFP#7-25 expressing the plastid marker FNR:eGFP (Schattat et al., 2011b) were used.

Plants were grown in long day conditions (16h day and 8h night) in green house chambers with controlled temperature and humidity. Temperature was approximately 23 °C during the day and 19 °C at night. Relative humidity was kept around 55 %.

### Bacterial strains and cultivation

*E. coli* Top10 cells were used for cloning and DNA propagation. Cells were cultivated at 37 °C in LB with the appropriate antibiotic selection. Agrobacterium strain GV3101 (pMP90) (Koncz & Schell, 1986) was grown in liquid or on solid YEB media containing rifampicin, gentamycin and either spectinomycin or carbenicillin, while *Xcv* strains were grown in NYG medium supplemented with rifampicin (30 °C for both). *Xanthomonas* strains utilized were: *Xcv* 85-10 (wild-type; Thieme *et al*., 2005), *Xcv ΔhrcN* (strain deficient in an ATPase required for typeIII secretion of effectors; Lorenz & Büttner, 2009 and *Xcv ΔxopQ* Adlung *et al*., 2016).

### Plasmids

For visualization of plastids and stromules a plastid organelle marker construct was created using the Modular Cloning Toolbox (Weber *et al*., 2011, Engler *et al*., 2014). The final construct consisted of the *35S* promotor (pICH51277), the chloroplast transit peptide of RUBISCO (pAGS-L0-#115 in backbone pICH41258), eGFP (pAGS-L0-#050 backbone pICH41264) and the NOS terminator (pICH41421), assembled in an Level 1 acceptor plasmid. The *XopQ:mOrange2* expression construct was described previously (Erickson et al., 2018; for more details see Methods S2).

### *Agrobacterium tumefaciens*-mediated transient expression

Plasmids were transformed into *A. tumefaciens* strain GV3101 (pMP90). For transient expression experiments, strains harboring the binary vectors were grown overnight in 5 ml YEB liquid cultures (with appropriate antibiotics), harvested by centrifugation and resuspended in agrobacterium infiltration medium (10 mM MgCl2; 5 mM MES pH 5.6; 0.15 µM Acetosyringone) with a final optical density (OD600nm) of 0.2. Bacteria harboring the plastid marker and the effector or the mORANGE2 control were mixed in a 1:1 ratio. Using a needless syringe, bacterial suspensions were inoculated into intercostal areas of the youngest fully expanded leaves of 5–6-week-old *N. benthamiana* plants (see Methods S3).

### *Xanthomonas campestris pv. vesicatoria* inoculations

*Xcv* NYG liquid cultures were centrifuged to harvest cells, bacteria was resuspended in 10 mM MgCl_2_, and suspensions adjusted to an OD600nm of 0.1. All three strains, as well as a buffer control, were then inoculated as described for *A. tumefaciens*. Plastids/stromules were observed at 2 days post-infiltration (dpi) using an epi-fluorescence microscope.

### Imaging hardware

For image acquisition, an epi-fluorescence microscope (AxioObserver Z1) setup from Zeiss (Jena, Germany) equipped with an X-Cite fluorescence light source and a MRm monochrome camera (Zeiss, Jena, Germany) was used. GFP fluorescence was recorded using a 38 HE filter cube (Carl Zeiss AG, Jena, Germany). mORANGE2 fluorescence was recorded utilizing the 43 HE filter cube (Carl Zeiss AG. Jena, Germany). The microscope manufacturer’s software (ZenBlue, Zeiss, Germany) controlled image acquisition. All images were captured using a 40x / 0.75 NA EC PLAN NEOFLUAR lens.

### Imaging procedures and image processing

For quantification of stromule frequencies, a single leaf disc of each infiltration spot was harvested using a cork borer. Leaf discs were vacuum-infiltrated, mounted on glass slides and three independent z-stacks of the lower epidermis were collected in transmitted light, eGFP and mORANGE2 channels. In order to obtain 2D extended depth of field images for quantification, single images of the z-series of each channel were first exported into separate file folders and subsequently combined into single images using software and procedures described in Schattat & Klösgen, 2009 (total of 3 images per disc).

For quantification of stromule frequencies (SF%) we measured the proportion of plastids with at least one stromule (Erickson et al., 2014). To facilitate the faster quantification of stromule and plastid counts in *N. benthamiana* tissues, we expanded on the previously published MTBCellCounter (Franke *et al*., 2015) via a ridge detection-based stromule detection algorithm (Möller & Schattat, 2019). The extended MTBCellCounter allows for the detection of plastid bodies as described in Franke *et al*., 2015 and identifies subsequently plastids with stromules.

The plastid nucleus association index (PNAI) was described previously (Erickson *et al*., 2014), and represents the absolute number of plastids in close association with a given nucleus. PNAI was evaluated in the 2D projected images (see image processing). Nuclei were counted as nucleus associated when either the plastid body touched, overlapped with, or was within a distance of 4µm from the nucleus. 4µm corresponds to the average epidermis plastid diameter.

For sample sizes and details on statistical analysis of SF% and PNAI see Supplemental materials Methods S4 Notes S1.

### Naming conventions

For conventions used to name mutants, genes, proteins and artificial DNA constructs see Methods S5.

## Supporting information

supplemental material stats

## Acknowledgments

All *Xcv* strains were kindly provided by Prof. Ulla Bonas, Martin-Luther-Universität Halle-Wittenberg, Germany. Dr. Sebastian Schornack, TSL, Cambridge, UK for advice on the manuscript. This study was funded by the Deutsche Forschungsgemeinschaft (DFG, German Research Foundation) – 400681449/GRK2498, STU642/1-1 and SFB-1403–414786233 as well as Martin-Luther-University core funding.

