## supplemental material tables for "XopQ induced stromule formation in *Nicotiana benthamiana* is causally linked to ETI signaling and depends on ADR1 and NRG1"

### Supplemental materials – tables

**Table S1 Values used for bar plots in Fig. S2.** n = total number of plants used for respective treatment and genotype; SF% Mean = arithmetic average of stromule frequency values expressed in %; C.I. of Mean 95% lower = absolute value of the lower 95% confidence interval of the presented mean; C.I. of Mean 95% upper = absolute value of the upper 95% confidence interval of the presented mean; difference C.I. Mean 95% lower = difference of mean and the absolute value of the lower 95% confidence interval; difference C.I. Mean 95% upper = difference of the absolute value of the 95% upper confidence interval and the mean; the difference values have been used to blot the whiskers in the bar blots.

| | n | SF% Mean | C.I. of Mean 95% lower | C.I. of Mean 95% upper | difference C.I. Mean 95% lower | difference C.I. Mean 95% upper |
|---|---|---|---|---|---|---|
| None infiltrated | 16 | 2 | 1.4 | 2.6 | 0.5 | 0.6 |
| AIM – buffer control | 22 | 5.7 | 3.9 | 7.7 | 1.7 | 2 |
| mORANGE2 | 23 | 19.7 | 15.4 | 24.4 | 4.3 | 4.7 |

**Table S2 Summary of values used for stromule frequency bar blots in the main manuscript**. Values used for drawing bar blots in Fig. 2a, 3a, 4a, 5a, 6a and 7a; n = total number of plants used for respective treatment and genotype; SF% Mean = arithmetic average of stromule frequency values expressed in %; C.I. of Mean 95% lower = absolute value of the lower 95% confidence interval of the presented mean; C.I. of Mean 95% upper = absolute value of the upper 95% confidence interval of the presented mean; difference C.I. Mean 95% lower = difference of mean and the absolute value of the lower 95% confidence interval; difference C.I. Mean 95% upper = difference of the absolute value of the 95% upper confidence interval and the mean; the difference values have been used to blot the whiskers in the bar blots.

| | n | SF% Mean | C.I. of Mean 95% lower | C.I. of Mean 95% upper | difference C.I. Mean 95% lower | difference C.I. Mean 95% upper |
|---|---|---|---|---|---|---|
| Figure 2 | | | | | | |
| Mock | 4 | 3 | 1.7 | 4.7 | 1.3 | 1.7 |
| *Xcv 85-10ΔhrcN* | 4 | 3.8 | 1.7 | 6.6 | 2.1 | 2.8 |
| *Xcv 85-10ΔxopQ* | 4 | 3.9 | 2.7 | 5.4 | 1.2 | 1.5 |
| *Xcv 85-10* | 4 | 32.3 | 24.8 | 40.2 | 7.45 | 8 |
| Figure 3 | | | | | | |
| *roq1-3* / mORANGE2 | 13 | 30 | 26.2 | 34 | 3.8 | 3.9 |
| *roq1-3* / XopQ | 13 | 27.1 | 20.3 | 34.4 | 6.8 | 7.3 |
| *roq1-4* / mORANGE2 | 13 | 23.4 | 17.8 | 29.5 | 5.6 | 6.1 |
| *roq1-4* / XopQ: mORANGE2 | 13 | 22.8 | 19.5 | 26.3 | 3.3 | 3.5 |
| WT / mORANGE2 | 13 | 25 | 18.7 | 32 | 6.3 | 6.9 |
| WT / XopQ: mORANGE2 | 13 | 55.1 | 44 | 65.9 | 11.1 | 10.8 |
| Figure 4 | | | | | | |
| *eds1a-1* / mORANGE2 | 11 | 12.5 | 10.2 | 15.1 | 2.4 | 2.6 |
| *eds1a-1* / XopQ | 11 | 8.8 | 10.8 | 7.1 | 1.7 | 1.9 |
| WT / mORANGE2 | 11 | 11.4 | 14.7 | 8.4 | 2.9 | 3.3 |
| WT / XopQ: mORANGE2 | 11 | 47.9 | 51.9 | 43.8 | 4 | 4 |
| Figure 5 | | | | | | |
| *nrg1-4* / mORANGE2 | 15 | 22.8 | 19.8 | 26 | 3 | 3.2 |
| *nrg1-4* / XopQ: mORANGE2 | 15 | 43.7 | 41.4 | 46 | 2.3 | 2.3 |
| *nrg1-5* / mORANGE2 | 15 | 24.6 | 20.8 | 28.6 | 3.8 | 4 |
| *nrg1-5* / XopQ: mORANGE2 | 15 | 47.1 | 42.4 | 51.7 | 4.6 | 4.7 |
| WT / mORANGE2 | 15 | 28.1 | 23.1 | 33.5 | 5 | 5.3 |
| WT / XopQ: mORANGE2 | 15 | 63.7 | 57.4 | 69.8 | 6.3 | 6.1 |
| Figure 6 | | | | | | |
| *adr1_nrg1* / mORANGE2 | 15 | 27.8 | 23.6 | 32.3 | 4.3 | 4.5 |
| *adr1_nrg1* / XopQ: mORANGE2 | 15 | 28.8 | 24.9 | 32.9 | 3.9 | 4.1 |
| WT / mORANGE2 | 9 | 32.5 | 26.8 | 38.5 | 5.7 | 5.9 |
| WT / XopQ: mORANGE2 | 9 | 54.9 | 46 | 63.7 | 8.9 | 8.8 |

**Table S3 PNAI summary of values used for box blots in the main manuscript.** Values displayed in the PNAI box blots of Fig. 8a. Infiltration = constructs co-infiltrated; plant line = genetic background used for experiment; n = number of nuclei included in the analysis. Lower numbers for X*opQ:mOrange2*-related experiments result from lower expression levels and therefore lower numbers of clearly defined nuclei. Median PNAI = median of the plastid nucleus association values; 25% = value for the 25 percentile; 75% = value of the 75 percentile.

| Expressed constructs | plant line | n | median PNAI | 25% | 75% |
|---|---|---|---|---|---|
| *mOrange2 + SSU:eGFP* | wild-type | 464 | 4 | 4 | 6 |
| *XopQ:mOrange2 + SSU:eGFP* | wild-type | 297 | 6 | 5.75 | 7 |
| *mOrange2 + SSU:eGFP* | roq1-3 | 95 | 5 | 4 | 6 |
| *XopQ:mOrange2 + SSU:eGFP* | roq1-3 | 61 | 7 | 6 | 9 |
| *mOrange2 + SSU:eGFP* | eds1a-1 | 90 | 5 | 3 | 7 |
| *XopQ:mOrange2 + SSU:eGFP* | eds1a-1 | 66 | 8 | 5 | 10 |
| *mOrange2 + SSU:eGFP* | nrg1-4 | 126 | 4 | 3 | 5 |
| *XopQ:mOrange2 + SSU:eGFP* | nrg1-4 | 113 | 6 | 5 | 7 |
| *mOrange2 + SSU:eGFP* | nrg1_adr1 | 249 | 5 | 3 | 6 |
| *XopQ:mOrange2 + SSU:eGFP* | nrg1_adr1 | 102 | 8 | 6 | 10.25 |
