## supplemental material stats for "XopQ induced stromule formation in *Nicotiana benthamiana* is causally linked to ETI signaling and depends on ADR1 and NRG1"

**Supplemental materials – statistics**

**Notes S1 Details of performed statistical tests for plots shown in the main manuscript.**

**Fig. 2 SF% *Xcv***

| Group | N | Missing | Median | 25% | 75% |
|---|---|---|---|---|---|
| XopQ | 12 | 0 | 0.204 | 0.157 | 0.234 |
| moc | 12 | 0 | 0.169 | 0.148 | 0.224 |

Mann-Whitney U Statistic= 57.000

T = 165.000  n(small)= 12  n(big)= 12  (P = 0.403)

The difference in the median values between the two groups is not great enough to exclude the possibility that the difference is due to random sampling variability; there is not a statistically significant difference (P = 0.403)

| Group | N | Missing | Median | 25% | 75% |
|---|---|---|---|---|---|
| moc | 12 | 0 | 0.169 | 0.148 | 0.224 |
| hrcN | 12 | 0 | 0.168 | 0.117 | 0.283 |

Mann-Whitney U Statistic= 70.000

T = 152.000  n(small)= 12  n(big)= 12  (P = 0.931)

The difference in the median values between the two groups is not great enough to exclude the possibility that the difference is due to random sampling variability; there is not a statistically significant difference (P = 0.931)

| Group | N | Missing | Median | 25% | 75% |
|---|---|---|---|---|---|
| moc | 12 | 0 | 0.169 | 0.148 | 0.224 |
| xcv | 12 | 0 | 0.605 | 0.512 | 0.680 |

Mann-Whitney U Statistic= 0.000

T = 78.000  n(small)= 12  n(big)= 12  (P = <0.001)

The difference in the median values between the two groups is greater than would be expected by chance; there is a statistically significant difference  (P = <0.001)

| Group | N | Missing | Median | 25% | 75% |
|---|---|---|---|---|---|
| XopQ | 12 | 0 | 0.204 | 0.157 | 0.234 |
| hrcN | 12 | 0 | 0.168 | 0.117 | 0.283 |

Mann-Whitney U Statistic= 60.000

T = 162.000  n(small)= 12  n(big)= 12  (P = 0.507)

The difference in the median values between the two groups is not great enough to exclude the possibility that the difference is due to random sampling variability; there is not a statistically significant difference (P = 0.507)

**Fig. 3 SF% *roq1***

| Group | N | Missing | Median | 25% | 75% |
|---|---|---|---|---|---|
| Roq1-3 mOrange | 13 | 0 | 0.602 | 0.511 | 0.624 |
| Roq1-3 XopQ | 13 | 0 | 0.577 | 0.509 | 0.635 |

Mann-Whitney U Statistic= 78.500

T = 181.500  n(small)= 13  n(big)= 13  (P = 0.778)

The difference in the median values between the two groups is not great enough to exclude the possibility that the difference is due to random sampling variability; there is not a statistically significant difference (P = 0.778)

| Group | N | Missing | Median | 25% | 75% |
|---|---|---|---|---|---|
| Roq1-4 mOrange | 13 | 0 | 0.542 | 0.438 | 0.592 |
| Roq1-4 XopQ | 13 | 0 | 0.515 | 0.451 | 0.533 |

Mann-Whitney U Statistic= 71.500

T = 188.500  n(small)= 13  n(big)= 13  (P = 0.521)

The difference in the median values between the two groups is not great enough to exclude the possibility that the difference is due to random sampling variability; there is not a statistically significant difference (P = 0.521)

| Group | N | Missing | Median | 25% | 75% |
|---|---|---|---|---|---|
| WT mOrange | 13 | 0 | 0.492 | 0.459 | 0.604 |
| WT XopQ | 13 | 0 | 0.915 | 0.655 | 0.981 |

Mann-Whitney U Statistic= 14.500

T = 105.500  n(small)= 13  n(big)= 13  (P = <0.001)

The difference in the median values between the two groups is greater than would be expected by chance; there is a statistically significant difference  (P = <0.001)

| Group | N | Missing | Median | 25% | 75% |
|---|---|---|---|---|---|
| WT mOrange | 13 | 0 | 0.492 | 0.459 | 0.604 |
| Roq1-3 mOrange | 13 | 0 | 0.602 | 0.511 | 0.624 |

Mann-Whitney U Statistic= 52.500

T = 143.500  n(small)= 13  n(big)= 13  (P = 0.106)

The difference in the median values between the two groups is not great enough to exclude the possibility that the difference is due to random sampling variability; there is not a statistically significant difference (P = 0.106)

| Group | N | Missing | Median | 25% | 75% |
|---|---|---|---|---|---|
| WT mOrange | 13 | 0 | 0.492 | 0.459 | 0.604 |
| Roq1-4 mOrange | 13 | 0 | 0.542 | 0.438 | 0.592 |

Mann-Whitney U Statistic= 82.500

T = 177.500  n(small)= 13  n(big)= 13  (P = 0.939)

The difference in the median values between the two groups is not great enough to exclude the possibility that the difference is due to random sampling variability; there is not a statistically significant difference (P = 0.939)

**Fig. 4 SF% *eds1a-1***

| Group | N | Missing | Median | 25% | 75% |
|---|---|---|---|---|---|
| WT mOrange2 | 11 | 0 | 0.359 | 0.271 | 0.457 |
| WT XopQ | 11 | 0 | 0.771 | 0.707 | 0.826 |

Mann-Whitney U Statistic= 0.000

T = 66.000 n(small)= 11 n(big)= 11 (P = <0.001)

The difference in the median values between the two groups is greater than would be expected by chance; there is a statistically significant difference (P = <0.001)

| Group | N | Missing | Median | 25% | 75% |
|---|---|---|---|---|---|
| eds1 mOrange2 | 11 | 0 | 0.381 | 0.271 | 0.435 |
| eds1 XopQ | 11 | 0 | 0.286 | 0.247 | 0.399 |

Mann-Whitney U Statistic= 43.000

T = 144.000 n(small)= 11 n(big)= 11 (P = 0.264)

The difference in the median values between the two groups is not great enough to exclude the possibility that the difference is due to random sampling variability; there is not a statistically significant difference (P = 0.264)

| Group | N | Missing | Median | 25% | 75% |
|---|---|---|---|---|---|
| eds1 mOrange2 | 11 | 0 | 0.381 | 0.271 | 0.435 |
| WT mOrange2 | 11 | 0 | 0.359 | 0.271 | 0.457 |

Mann-Whitney U Statistic= 56.000

T = 131.000 n(small)= 11 n(big)= 11 (P = 0.793)

The difference in the median values between the two groups is not great enough to exclude the possibility that the difference is due to random sampling variability; there is not a statistically significant difference (P = 0.793)

**Fig. 5 SF% *nrg1***

| Group | N | Missing | Median | 25% | 75% |
|---|---|---|---|---|---|
| WT mOrange2 | 15 | 0 | 0.529 | 0.484 | 0.627 |
| WT XopQ | 15 | 0 | 0.934 | 0.842 | 1.010 |

Mann-Whitney U Statistic= 2.000

T = 122.000 n(small)= 15 n(big)= 15 (P = <0.001)

The difference in the median values between the two groups is greater than would be expected by chance; there is a statistically significant difference (P = <0.001)

| Group | N | Missing | Median | 25% | 75% |
|---|---|---|---|---|---|

| | | | | | |
|---|---|---|---|---|---|
| nrg1-5 mOrange2 | 15 | 0 | 0.507 | 0.473 | 0.589 |
| nrg1-5 XopQ | 15 | 0 | 0.720 | 0.691 | 0.784 |

Mann-Whitney U Statistic= 0.000

T = 120.000 n(small)= 15 n(big)= 15 (P = <0.001)

The difference in the median values between the two groups is greater than would be expected by chance; there is a statistically significant difference (P = <0.001)

| Group | N | Missing | Median | 25% | 75% |
|---|---|---|---|---|---|
| nrg1-4 mOrange2 | 15 | 0 | 0.512 | 0.458 | 0.541 |
| nrg1-4 XopQ | 15 | 0 | 0.718 | 0.696 | 0.754 |

Mann-Whitney U Statistic= 0.000

T = 120.000 n(small)= 15 n(big)= 15 (P = <0.001)

The difference in the median values between the two groups is greater than would be expected by chance; there is a statistically significant difference (P = <0.001)

| Group | N | Missing | Median | 25% | 75% |
|---|---|---|---|---|---|
| WT mOrange2 | 15 | 0 | 0.529 | 0.484 | 0.627 |
| nrg1-4mOrange2 | 15 | 0 | 0.512 | 0.458 | 0.541 |

Mann-Whitney U Statistic= 80.000

T = 265.000 n(small)= 15 n(big)= 15 (P = 0.184)

The difference in the median values between the two groups is not great enough to exclude the possibility that the difference is due to random sampling variability; there is not a statistically significant difference (P = 0.184)

| Group | N | Missing | Median | 25% | 75% |
|---|---|---|---|---|---|
| WT mOrange2 | 15 | 0 | 0.529 | 0.484 | 0.627 |
| nrg1-5 mOrange2 | 15 | 0 | 0.507 | 0.473 | 0.589 |

Mann-Whitney U Statistic= 90.500

T = 254.500 n(small)= 15 n(big)= 15 (P = 0.372)

The difference in the median values between the two groups is not great enough to exclude the possibility that the difference is due to random sampling variability; there is not a statistically significant difference (P = 0.372)

| Group | N | Missing | Median | 25% | 75% |
|---|---|---|---|---|---|
| WT XopQ | 15 | 0 | 0.934 | 0.842 | 1.010 |
| nrg1-4 XopQ | 15 | 0 | 0.718 | 0.696 | 0.754 |

Mann-Whitney U Statistic= 15.000

T = 330.000 n(small)= 15 n(big)= 15 (P = <0.001)

The difference in the median values between the two groups is greater than would be expected by chance; there is a statistically significant difference (P = <0.001)

| Group | N | Missing | Median | 25% | 75% |
|---|---|---|---|---|---|
| WT XopQ | 15 | 0 | 0.934 | 0.842 | 1.010 |
| nrg1-5 XopQ | 15 | 0 | 0.720 | 0.691 | 0.784 |

Mann-Whitney U Statistic= 28.000

T = 317.000 n(small)= 15 n(big)= 15 (P = <0.001)

The difference in the median values between the two groups is greater than would be expected by chance; there is a statistically significant difference (P = <0.001)

**Fig. 6 SF% *adr1_nrg1***

| Group | N | Missing | Median | 25% | 75% |
|---|---|---|---|---|---|
| WT_mOrange | 9 | 0 | 0.560 | 0.550 | 0.660 |
| WT_XopQ | 9 | 0 | 0.854 | 0.786 | 0.910 |

Mann-Whitney U Statistic= 6.000

T = 51.000 n(small)= 9 n(big)= 9 (P = 0.003)

The difference in the median values between the two groups is greater than would be expected by chance; there is a statistically significant difference (P = 0.003)

| Group | N | Missing | Median | 25% | 75% |
|---|---|---|---|---|---|
| Adr1/Nrg1_mOrange | 15 | 0 | 0.539 | 0.490 | 0.578 |
| Adr1/Nrg1_XopQ | 15 | 0 | 0.549 | 0.514 | 0.625 |

Mann-Whitney U Statistic= 99.000

T = 219.000 n(small)= 15 n(big)= 15 (P = 0.590)

The difference in the median values between the two groups is not great enough to exclude the possibility that the difference is due to random sampling variability; there is not a statistically significant difference (P = 0.590)

| Group | N | Missing | Median | 25% | 75% |
|---|---|---|---|---|---|
| WT_XopQ | 9 | 0 | 0.854 | 0.786 | 0.910 |
| Adr1/Nrg1_XopQ | 15 | 0 | 0.549 | 0.514 | 0.625 |

Mann-Whitney U Statistic= 6.000

T = 174.000 n(small)= 9 n(big)= 15 (P = <0.001)

The difference in the median values between the two groups is greater than would be expected by chance; there is a statistically significant difference (P = <0.001)

| Group | N | Missing | Median | 25% | 75% |
|---|---|---|---|---|---|
| WT_mOrange | 9 | 0 | 0.560 | 0.550 | 0.660 |
| Adr1/Nrg1_mOrange | 15 | 0 | 0.539 | 0.490 | 0.578 |

Mann-Whitney U Statistic= 43.000

T = 137.000 n(small)= 9 n(big)= 15 (P = 0.152)

The difference in the median values between the two groups is not great enough to exclude the possibility that the difference is due to random sampling variability; there is not a statistically significant difference (P = 0.152)

**Fig. 7 PNAI**

| Group | N | Missing | Median | 25% | 75% |
|---|---|---|---|---|---|
| Roq1-3 mOrange | 95 | 0 | 5.000 | 4.000 | 6.000 |
| Roq-1-3 XopQ | 61 | 0 | 7.000 | 5.500 | 9.000 |

Mann-Whitney U Statistic= 1435.500

T = 6250.500 n(small)= 61 n(big)= 95 (P = <0.001)

The difference in the median values between the two groups is greater than would be expected by chance; there is a statistically significant difference (P = <0.001)

| Group | N | Missing | Median | 25% | 75% |
|---|---|---|---|---|---|
| eds1 mOrange | 90 | 0 | 5.000 | 4.000 | 7.000 |
| eds1 XopQ | 66 | 0 | 8.000 | 6.000 | 10.000 |

Mann-Whitney U Statistic= 1279.000

T = 6872.000 n(small)= 66 n(big)= 90 (P = <0.001)

The difference in the median values between the two groups is greater than would be expected by chance; there is a statistically significant difference (P = <0.001)

| Group | N | Missing | Median | 25% | 75% |
|---|---|---|---|---|---|
| nrg1-4 mOrange2 | 126 | 0 | 4.000 | 3.000 | 5.000 |
| nrg1-4 XopQ | 113 | 0 | 6.000 | 5.000 | 7.000 |

Mann-Whitney U Statistic= 3170.000

T = 17509.000 n(small)= 113 n(big)= 126 (P = <0.001)

The difference in the median values between the two groups is greater than would be expected by chance; there is a statistically significant difference (P = <0.001)

| Group | N | Missing | Median | 25% | 75% |
|---|---|---|---|---|---|
| adr1/nrg1 mOrange2 | 249 | 0 | 5.000 | 3.000 | 6.000 |
| adr1/nrg1 XopQ | 102 | 0 | 8.000 | 6.000 | 10.250 |

Mann-Whitney U Statistic= 4408.500

T = 26242.500 n(small)= 102 n(big)= 249 (P = <0.001)

The difference in the median values between the two groups is greater than would be expected by chance; there is a statistically significant difference (P = <0.001)

| Group | N | Missing | Median | 25% | 75% |
|---|---|---|---|---|---|
| WT mOrange all | 464 | 0 | 4.000 | 3.000 | 6.000 |
| WT XopQ all | 297 | 0 | 6.000 | 4.000 | 7.000 |

Mann-Whitney U Statistic= 40668.000

T = 141393.000  n(small)= 297  n(big)= 464  (P = <0.001)

The difference in the median values between the two groups is greater than would be expected by chance; there is a statistically significant difference  (P = <0.001)

| Group | N | Missing | Median | 25% | 75% |
|---|---|---|---|---|---|
| Roq-1-3 XopQ | 61 | 0 | 7.000 | 5.500 | 9.000 |
| WT XopQ all | 297 | 0 | 6.000 | 4.000 | 7.000 |

Mann-Whitney U Statistic= 6449.500

T = 13558.500  n(small)= 61  n(big)= 297  (P = <0.001)

The difference in the median values between the two groups is greater than would be expected by chance; there is a statistically significant difference  (P = <0.001)

| Group | N | Missing | Median | 25% | 75% |
|---|---|---|---|---|---|
| eds1 XopQ | 66 | 0 | 8.000 | 6.000 | 10.000 |
| WT XopQ all | 297 | 0 | 6.000 | 4.000 | 7.000 |

Mann-Whitney U Statistic= 5171.000

T = 16642.000  n(small)= 66  n(big)= 297  (P = <0.001)

The difference in the median values between the two groups is greater than would be expected by chance; there is a statistically significant difference  (P = <0.001)

| Group | N | Missing | Median | 25% | 75% |
|---|---|---|---|---|---|
| nrg1-4 XopQ | 113 | 0 | 6.000 | 5.000 | 7.000 |
| WT XopQ all | 297 | 0 | 6.000 | 4.000 | 7.000 |

Mann-Whitney U Statistic= 16525.500

T = 22966.500  n(small)= 113  n(big)= 297  (P = 0.810)

The difference in the median values between the two groups is not great enough to exclude the possibility that the difference is due to random sampling variability; there is not a statistically significant difference (P = 0.810)

| Group | N | Missing | Median | 25% | 75% |
|---|---|---|---|---|---|
| adr1/nrg1 XopQ | 102 | 0 | 8.000 | 6.000 | 10.250 |
| WT XopQ all | 297 | 0 | 6.000 | 4.000 | 7.000 |

Mann-Whitney U Statistic= 7892.500

T = 27654.500  n(small)= 102  n(big)= 297  (P = <0.001)

The difference in the median values between the two groups is greater than would be expected by chance; there is a statistically significant difference  (P = <0.001)

| Group | N | Missing | Median | 25% | 75% |
|---|---|---|---|---|---|
| Roq-1-3 XopQ | 61 | 0 | 7.000 | 5.500 | 9.000 |

| | | | | | |
|---|---|---|---|---|---|
| eds1 XopQ | 66 | 0 | 8.000 | 6.000 | 10.000 |

Mann-Whitney U Statistic= 1608.000

T = 3499.000  n(small)= 61  n(big)= 66  (P = 0.049)

The difference in the median values between the two groups is greater than would be expected by chance; there is a statistically significant difference  (P = 0.049)

| Group | N | Missing | Median | 25% | 75% |
|---|---|---|---|---|---|
| eds1 XopQ | 66 | 0 | 8.000 | 6.000 | 10.000 |
| nrg1-4 XopQ | 113 | 0 | 6.000 | 5.000 | 7.000 |

Mann-Whitney U Statistic= 1787.000

T = 7882.000  n(small)= 66  n(big)= 113  (P = <0.001)

The difference in the median values between the two groups is greater than would be expected by chance; there is a statistically significant difference  (P = <0.001)

| Group | N | Missing | Median | 25% | 75% |
|---|---|---|---|---|---|
| nrg1-4 XopQ | 113 | 0 | 6.000 | 5.000 | 7.000 |
| adr1/nrg1 XopQ | 102 | 0 | 8.000 | 6.000 | 10.250 |

Mann-Whitney U Statistic= 2757.000

T = 14022.000  n(small)= 102  n(big)= 113  (P = <0.001)

The difference in the median values between the two groups is greater than would be expected by chance; there is a statistically significant difference  (P = <0.001)

| Group | N | Missing | Median | 25% | 75% |
|---|---|---|---|---|---|
| Roq-1-3 XopQ | 61 | 0 | 7.000 | 5.500 | 9.000 |
| adr1/nrg1 XopQ | 102 | 0 | 8.000 | 6.000 | 10.250 |

Mann-Whitney U Statistic= 2360.500

T = 4251.500  n(small)= 61  n(big)= 102  (P = 0.010)

The difference in the median values between the two groups is greater than would be expected by chance; there is a statistically significant difference  (P = 0.010)

| Group | N | Missing | Median | 25% | 75% |
|---|---|---|---|---|---|
| eds1 XopQ | 66 | 0 | 8.000 | 6.000 | 10.000 |
| adr1/nrg1 XopQ | 102 | 0 | 8.000 | 6.000 | 10.250 |

Mann-Whitney U Statistic= 3138.500

T = 5349.500  n(small)= 66  n(big)= 102  (P = 0.458)

The difference in the median values between the two groups is not great enough to exclude the possibility that the difference is due to random sampling variability; there is not a statistically significant difference (P = 0.458)

**Fig. S3 SF% GV3101(pMP90)**

| Group | N | Missing | Median | 25% | 75% |
| --- | --- | --- | --- | --- | --- |
| GV mOrange | 23 | 0 | 0.484 | 0.379 | 0.579 |
| ni | 16 | 0 | 0.132 | 0.119 | 0.156 |

Mann-Whitney U Statistic= 2.000

T = 138.000  n(small)= 16  n(big)= 23  (P = <0.001)

The difference in the median values between the two groups is greater than would be expected by chance; there is a statistically significant difference  (P = <0.001)

| Group | N | Missing | Median | 25% | 75% |
| --- | --- | --- | --- | --- | --- |
| AIM | 22 | 0 | 0.233 | 0.170 | 0.298 |
| GV mOrange | 23 | 0 | 0.484 | 0.379 | 0.579 |

Mann-Whitney U Statistic= 46.000

T = 299.000  n(small)= 22  n(big)= 23  (P = <0.001)

The difference in the median values between the two groups is greater than would be expected by chance; there is a statistically significant difference  (P = <0.001)
