## supplemental material methods for "XopQ induced stromule formation in *Nicotiana benthamiana* is causally linked to ETI signaling and depends on ADR1 and NRG1"

### Supplemental materials - Methods

**Methods S1 Generation of *Nbnrg1_adr1* double mutant line.** (**A**) Scheme of pDGE365 used for generation of *Nb_nrg1 adr1* double mutant plants; based on pDGE311 (Stuttmann *et al.*, 2021). Guide RNAs were expressed under control of an *Arabidopsis* U6 promotor fragment. Target sites are depicted, color code corresponds to panel b. (**B**) Gene models of *nrg1* and *adr1* genes targeted for editing. *NRG1* contains +1 and -1 nt mutations in the line used in this study; the +1 mutation at target site 1 induces a STOP codon directly downstream. The +1 insertion at target site 4 in *adr1* (magenta) induces an early STOP 20 codons downstream, before target site 5 (grey).

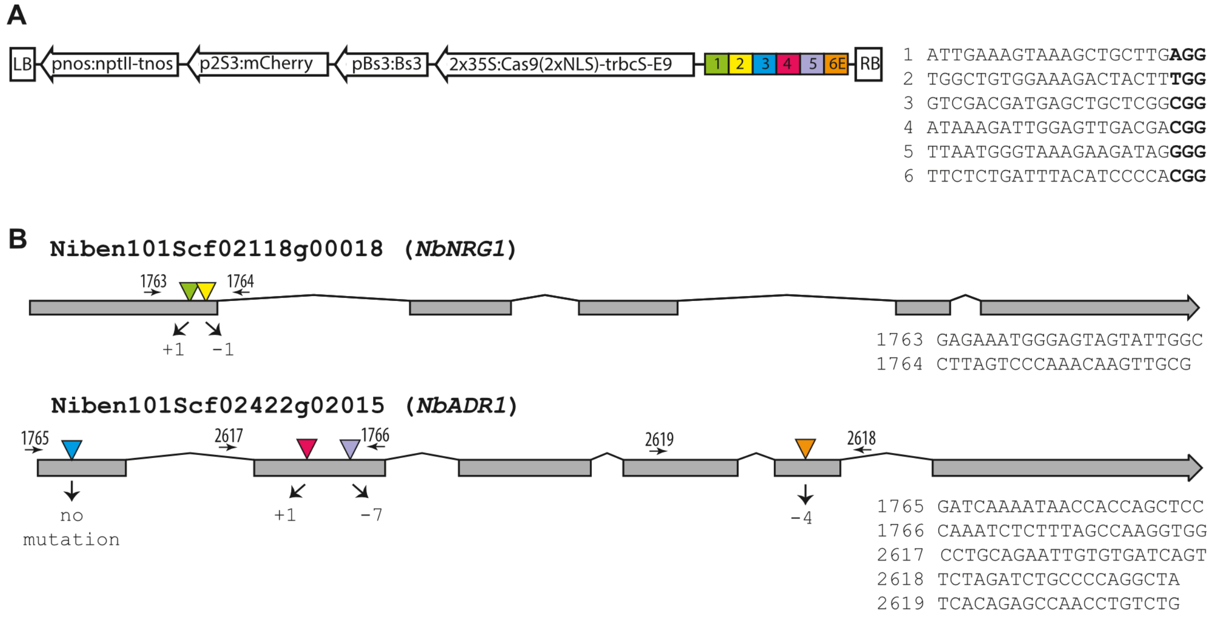

**Methods S2 Cloning of the plastid marker construct.** The promotor and terminator module were available in the plasmid collection of Engler et al. (2014). The transit peptide of RUBISCO from *Nicotiana benthamiana* was created as a SP module and eGFP as a CDS2* module (Engler *et al.*, 2014; Marillonnet & Werner, 2015). The following primers were used to clone the two modules: L0-SP-ssu-for = ttga aga caa aatg gct tcc tca gtt ctt tc; L0-SP-ssu-rev = ttg aag aca aac ctg agc tca aat cag gaa ggt atg; L0-CDS2*-eGFP-for = ttg aag aca aag gtg tga gca agg gcg agg; L0-CDS2*-eGFP-rev = ttg aag aca aaa gct tac ttg tac agc tcg tcc atg c. For the amplification of the transit peptide a plastid marker construct described in (Nelson *et al.*, 2007) was used as a template. eGFP was amplified from pICSL30006 (Engler *et al.*, 2014).

**Methods S3 Experimental procedure utilized for *A. tumefaciens* infiltration experiments.** Depiction of the experimental procedure for assessing stromule frequency in different mutant backgrounds. Bacteria mix = 1:1 mixes of *A. tumefaciens* of an $OD_{600nm}$ = 0.2.

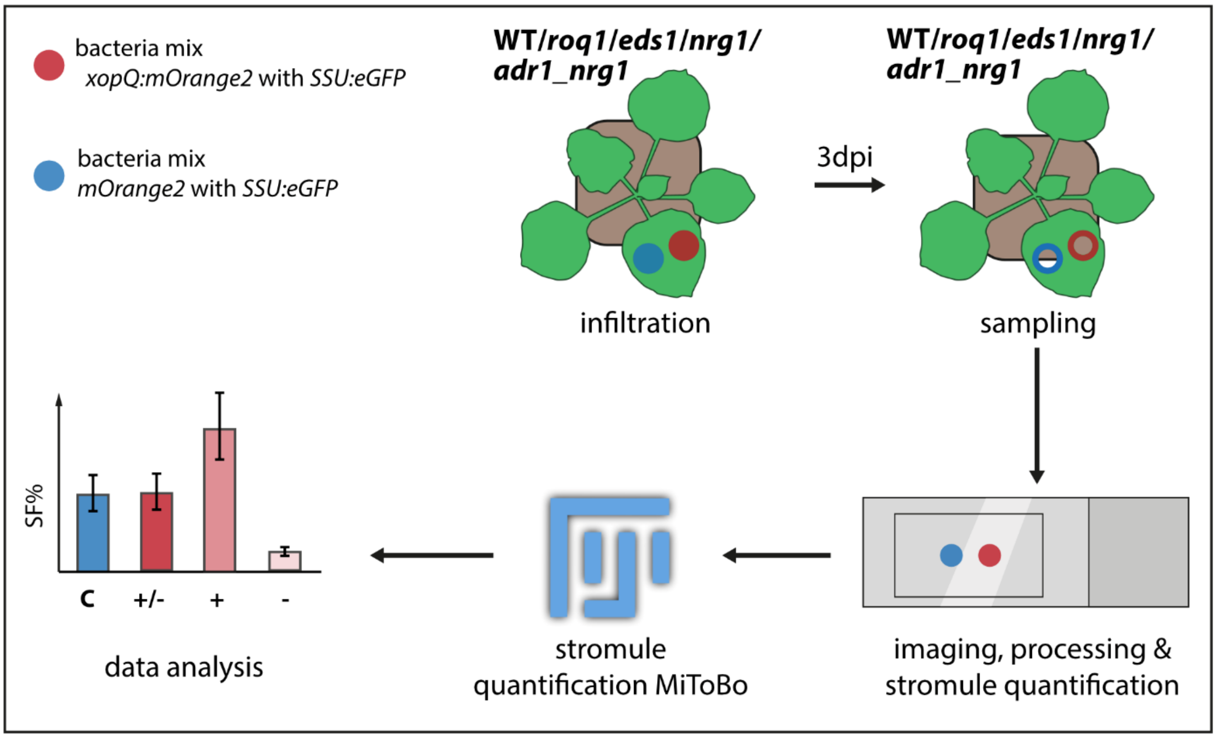

**Methods S4 Information on sample sizes and data analysis for stromule frequencies and PNAI values.** Stromule frequency measurements in WT, *roq1*, *eds1*, *nrg1* and *adr1_nrg1* mutant plant lines in response to *xopQ-mOrange2* and *mOrange2* expression were performed in 3 independently grown batches of 3-5 plants each. Stromule frequencies presented in Fig. 3, 4, 5 and 6 represent the average values of all experiments. Fig. 2 represents the average of 5 plants. For calculation of stromule frequencies in % (SF%) the number of plastids with one or more stromules was counted and divided by the total number of plastids. The resulting data was arcsin transformed, and statistical analysis was performed on the transformed data using SigmaPlot 12 (Systat Software GmbH, Erkrath, Germany). 95% confidence intervals and arithmetic averages were calculated and back-transformed data was represented in bar graphs (transformations completed using Microsoft Excel). For evaluating statistical significance between SF% values pairwise Mann-Whitney Rank Sum Tests were performed using SigmaPlot 12 (Systat Software GmbH, Erkrath, Germany). Test results can be found in Supporting Information – Statistics.

PNAI was measured with the help of the Fiji/ImageJ MTBCellclounterPlugIn (Franke *et al.*, 2015). For evaluating statistical significance between the PNAI of the various treatments Mann-Whitney Rank Sum Tests were performed using SigmaPlot 12 (Systat Software GmbH, Erkrath, Germany).

**Methods S5 Naming conventions.**

Genes

Names for plant genes are given in italic capital letters (e.g. *ROQ1*). Names of mutant alleles are printed in small italic letters (e.g. *roq1-3*). Bacterial genes are printed in small italic letters, with the exception of the letter designation which is capitalized (e.g. *xopQ* = the gene *Xanthomonas outer protein Q*). Genes which are missing in a given bacterial strain are indicated by the Greek letter delta (e.g. *ΔxopQ* = strain missing the *xopQ* gene). Cloned DNA sequences are handled like the respective genes. Fusion sites of DNA sequences are indicated by ":" (e.g. *FNR:eGFP; xopQ:mOrange2*).

Proteins

Plant proteins are not italicized and in all capital letters (e.g. ROQ1). Bacterial proteins are not italicized and start with a capital letter (e.g. XopQ = the protein Xanthomonas outer protein Q). Fluorescence proteins and genes are treated as plant genes and proteins when the name is abbreviated, such as *eGFP* (gene) or eGFP (Protein). Such protein names often consist of information about the species, the name of a color, descriptive words or oligomerization or other properties (this is indicated in small letters). In cases where the full name of the fluorescence protein is used, rather than the abbreviation, the first letter is capitalized (e.g. *mOrange2* = monomeric (property of protein=small) Orange fluorescence protein (full fluorescence protein name) 2 (numerical designation)). The protein is addressed in all capital not italicized letters except for the oligomerization descriptor (e.g. mORANGE2).
