## supplemental material figures for "XopQ induced stromule formation in *Nicotiana benthamiana* is causally linked to ETI signaling and depends on ADR1 and NRG1"

**Supplemental materials – Figures**

**Fig. S1 Full frames of stacked fluorescence images of a mock and *Xanthomonas campestris pv. vesicatoria ΔhrcN* infiltrated *Nicotiana benthamiana* plant**. Representative full-frame images used for stromule counts in *FNR:eGFP* transgenic plants in response to the mock treatment (**A**) and treatment with *Xcv hrcN* (**B**). Scale bar is 10 µm.

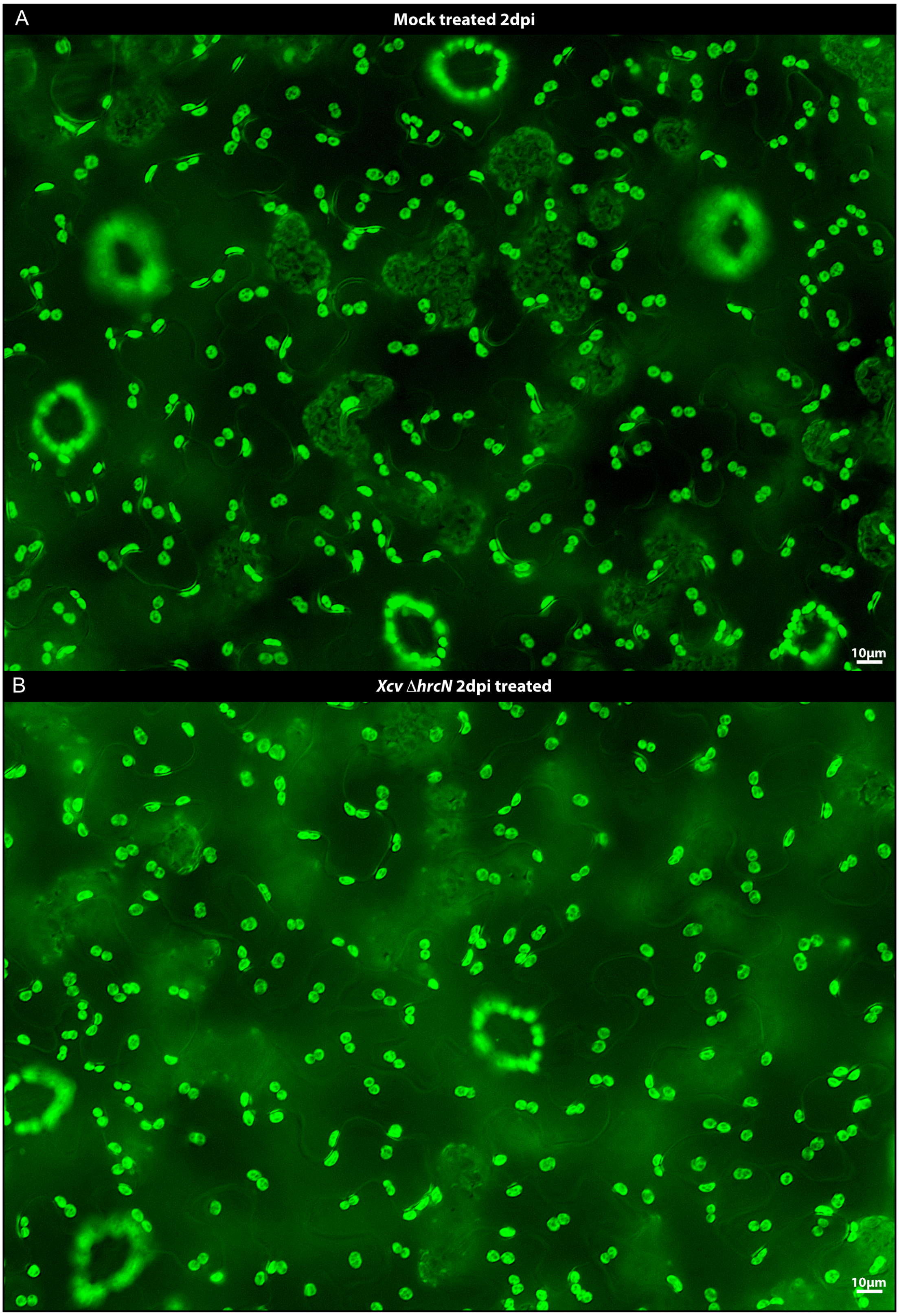

**Fig. S2 Full-frame of stacked fluorescence images of a *Xanthomonas campestris pv. vesicatoria 85-10* and *Xcv ΔxopQ* inoculated *Nicotiana benthamiana* plants**.

Representative full frame images used for stromule counts in *FNR:eGFP* transgenic plants in response to treatment with *Xcv 85-10* (A) and treatment with *Xcv ΔxopQ* (B).

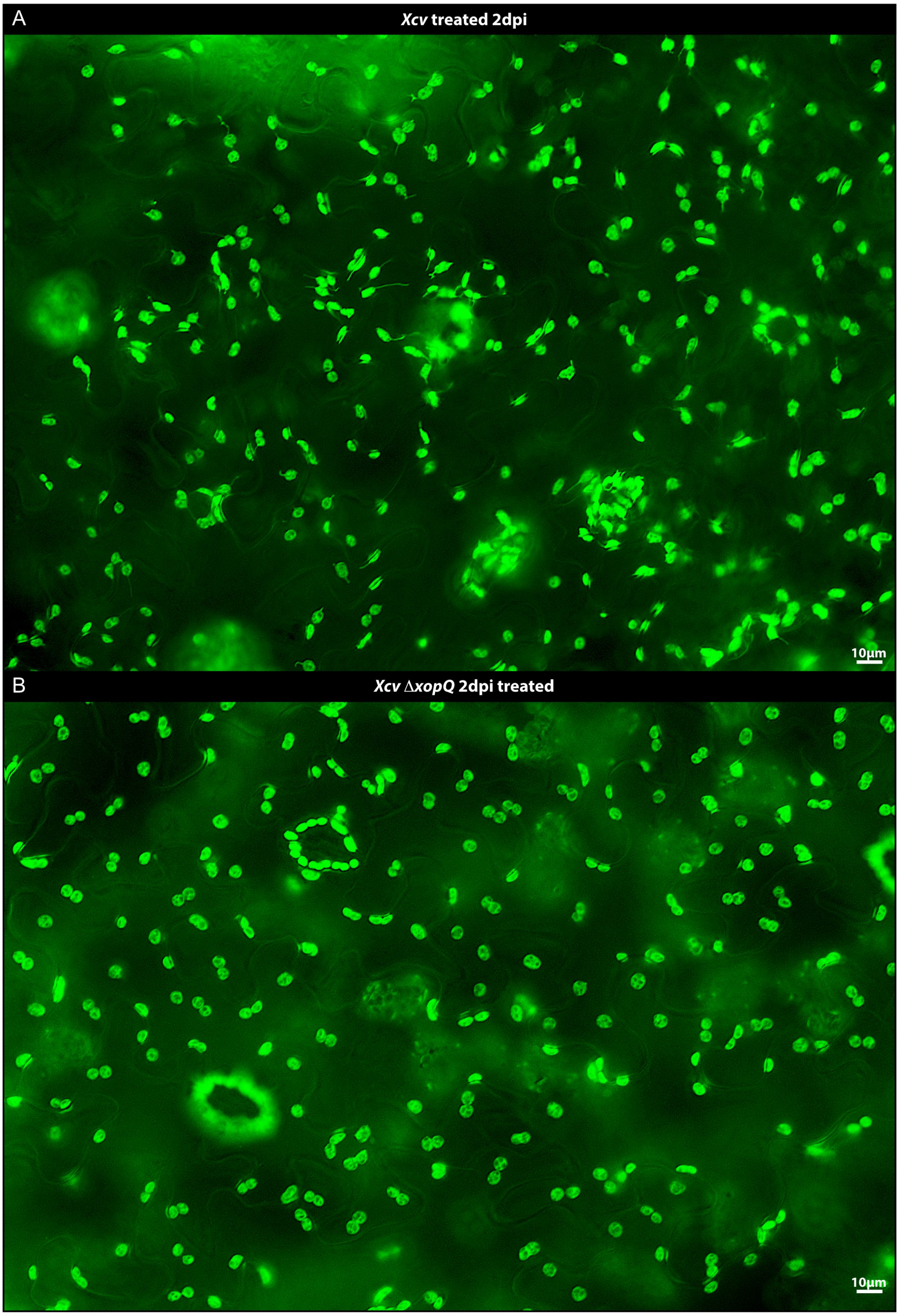

**Fig. S3 Moderate optical densities of GV3101 (pMP90) expressing *mOrange2* induce moderate stromule frequencies at 3 dpi**. (**A**) average stromule frequencies (SF%) in lower leaf epidermis cells of non-infiltrated (NI), agrobacterium infiltration medium (AIM)-infiltrated and GV3101 (pMP90)-infiltrated *N. benthamiana FNR:eGFP* plants. GV3101 (pMP90) mediates the expression of *mOrange2*. (**B**), (**C**) and (**D**) are representative sectors of images taken for stromule quantification. “n” = nucleus, arrow = stromule; scale bar corresponds to 10µm; green fluorescence originates from the *SSU:eGFP* plastid stroma marker (b, c and d) and the red fluorescence originates from the mORANGE2 fluorescence protein (**d**). For statistical values see Table S1.

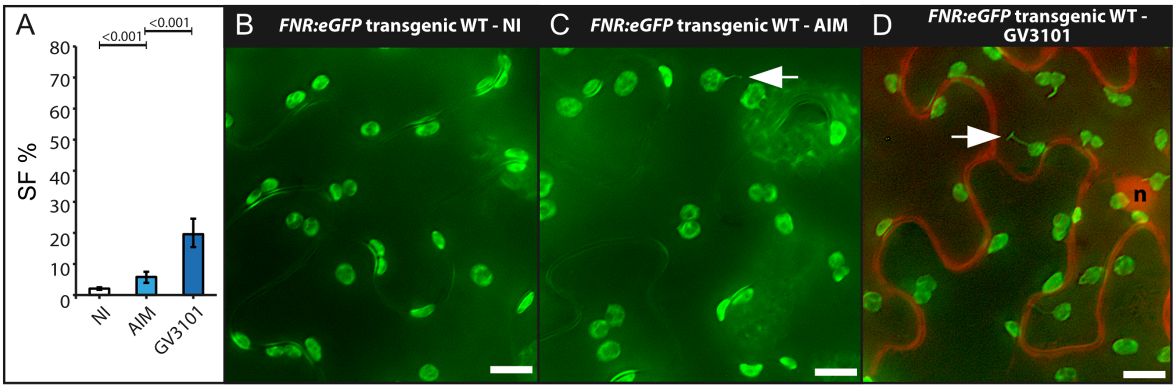

**Fig. S4 Full-frame stacked fluorescence images of an inoculated *Nicotiana benthamiana* wild-type plant.** Representative full frame images from the dataset used for stromule counts in WT in response to (**A**) *mOrange2+SSU:eGFP* and (**B**) *xopQ:mOrange2+SSU:eGFP*. Scale bars correspond to 10 µm.

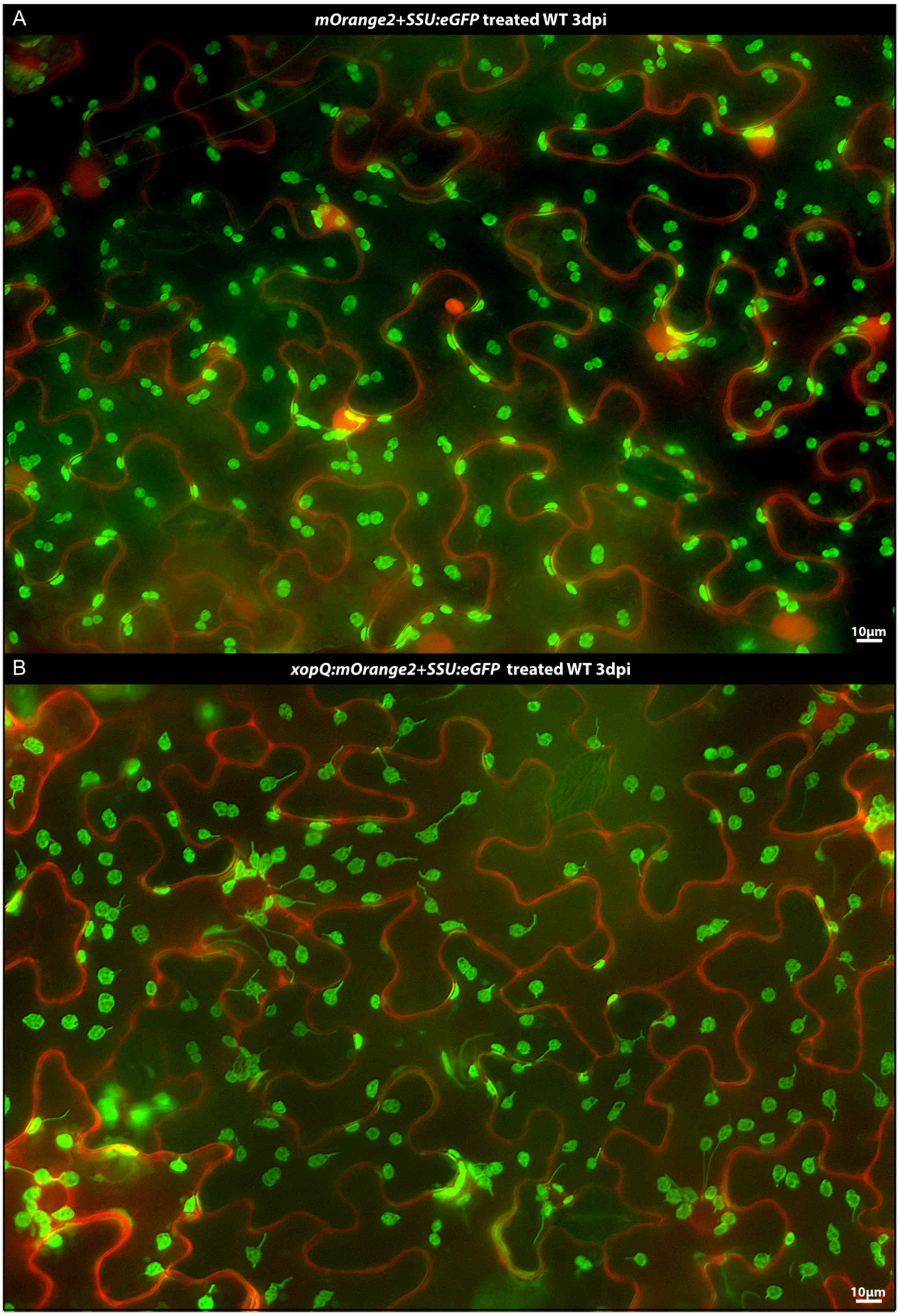

**Fig. S5 Full frame stacked fluorescence images of an inoculated *Nicotiana benthamiana roq1* plant.** Representative full-frame images used for stromule counts in *roq1-3* mutant in response to (**A**) *mOrange2+SSU:eGFP* and (**B**) *xopQ:mOrange2+SSU:eGFP*. Scale bars correspond to 10 µm.

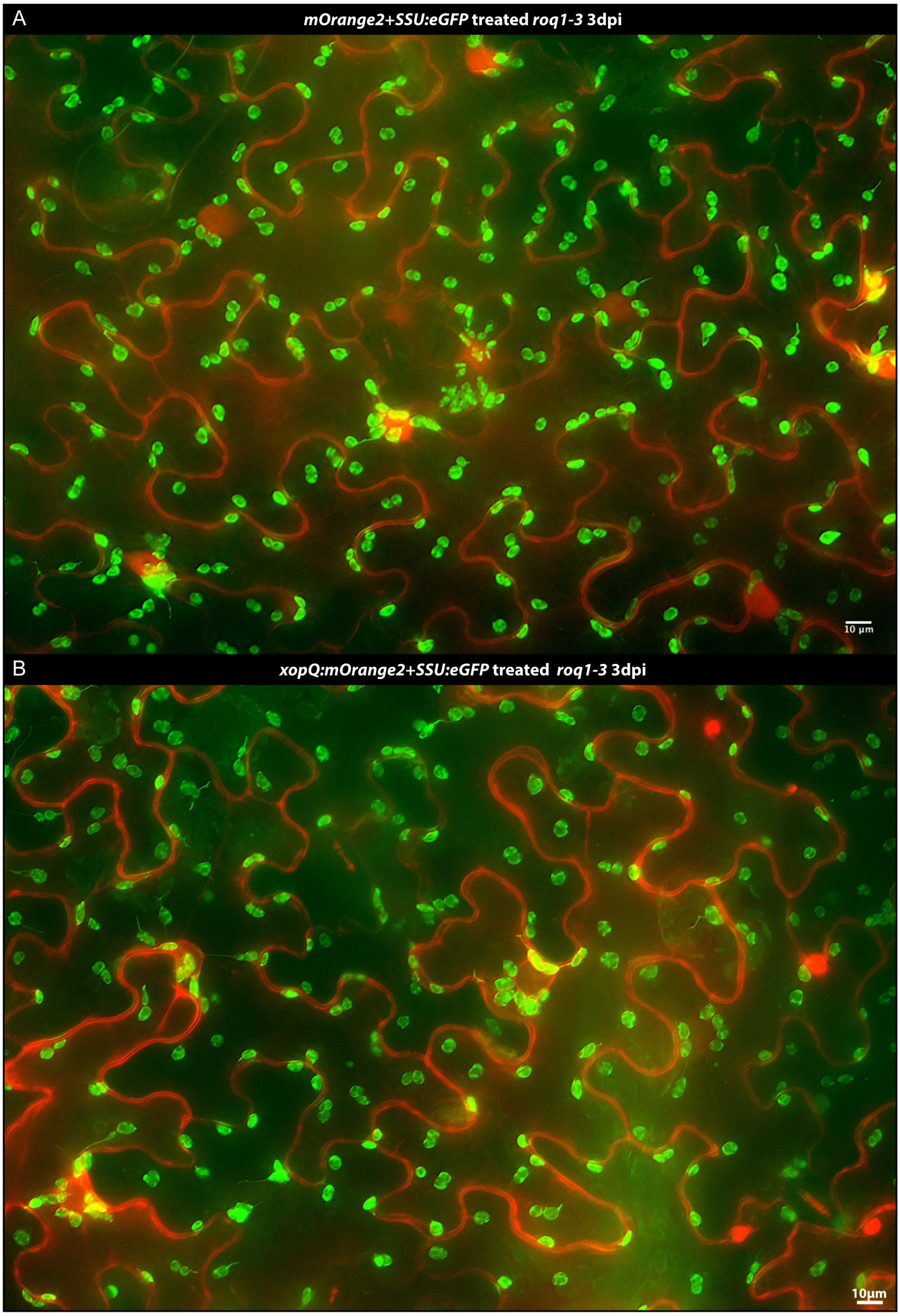

**Fig. S6 Full-frame stacked fluorescence images of an inoculated *Nicotiana benthamiana eds1* plant.** Representative full frame images from the dataset used for stromule counts in the *eds1a* mutant in response to (**A**) *mOrange2+SSU:eGFP* and (**B**) *xopQ:mOrange2+SSU:eGFP*. Scale bar corresponds to 10 µm.

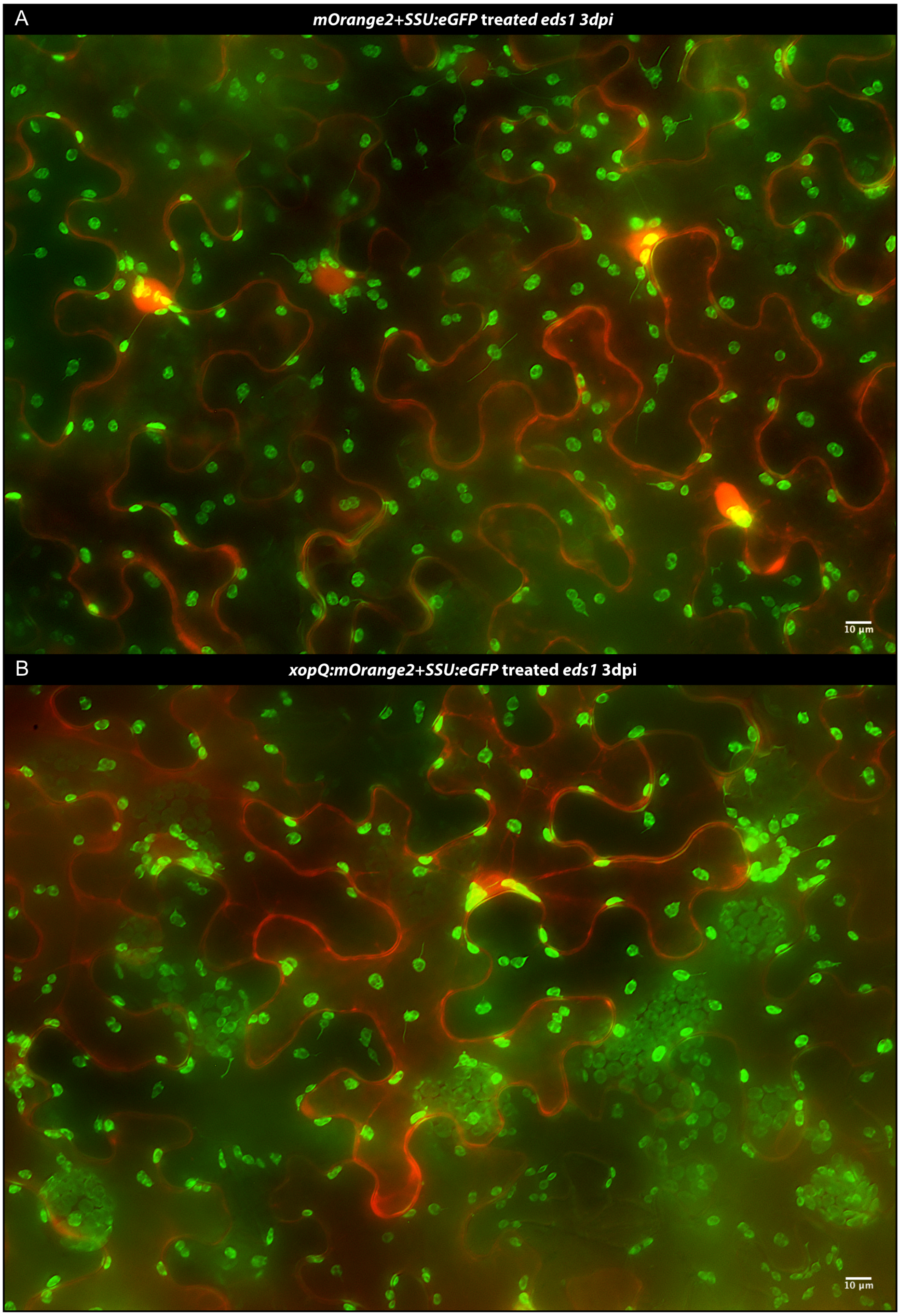

**Fig. S7 Full-frame stacked fluorescence images of an inoculated *Nicotiana benthamiana* *nrg1* plant.** Representative full-frame images used for stromule counts in the *nrg1-4* mutant in response to (**A**) *mOrange2+SSU:eGFP* and (**B**) *xopQ:mOrange2+SSU:eGFP*. Scale bar corresponds to 10 µm.

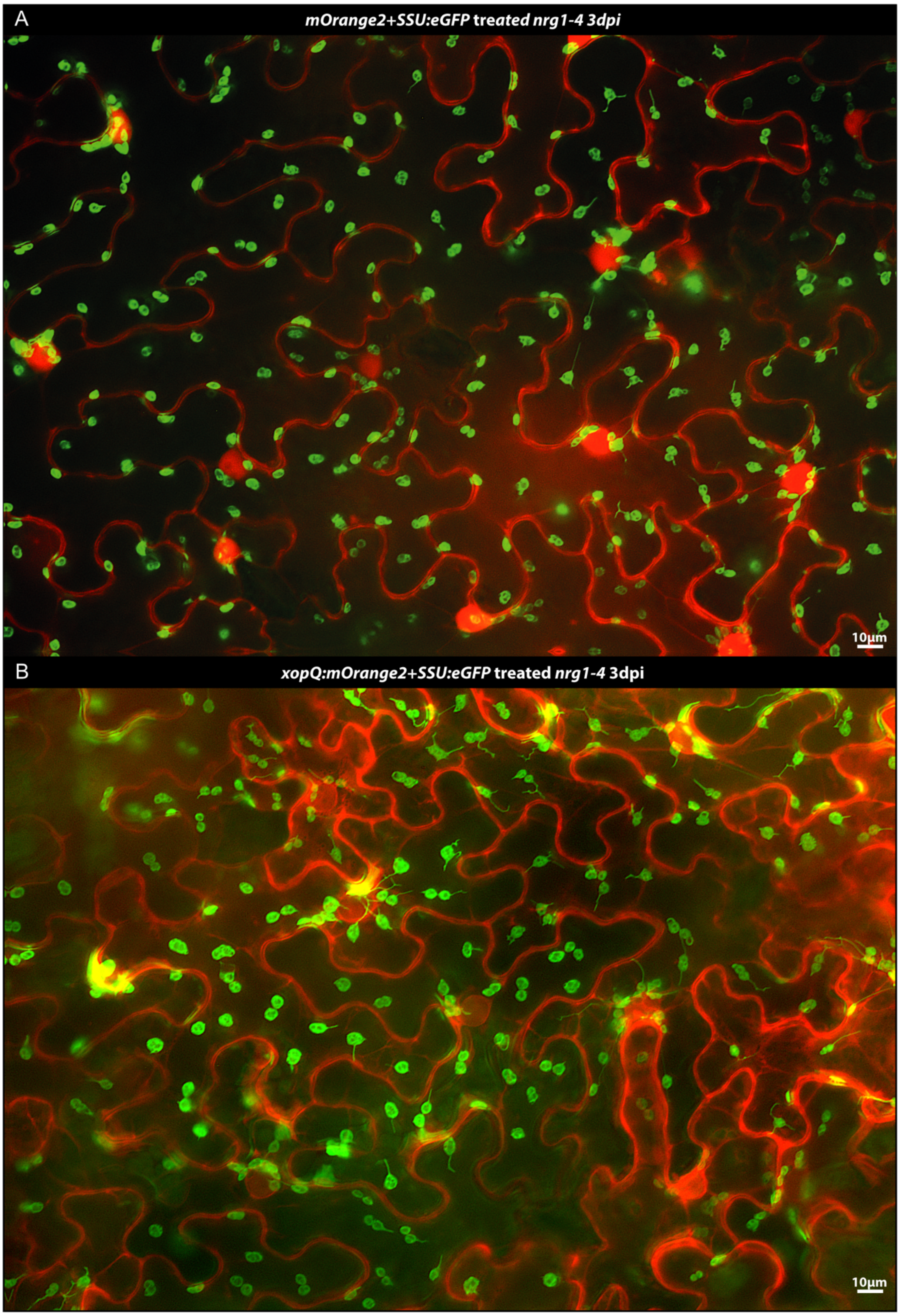

**Fig. S8 Full-frame stacked fluorescence images of an inoculated *Nicotiana benthamiana adr1_nrg1* plant.** Representative full-frame images used for stromule counts in the *adr1_nrg1* mutant in response to (**A**) *mOrange2+SSU:eGFP* and (**B**) *xopQ:mOrange2+SSU:eGFP*. Scale bar corresponds to 10 µm.

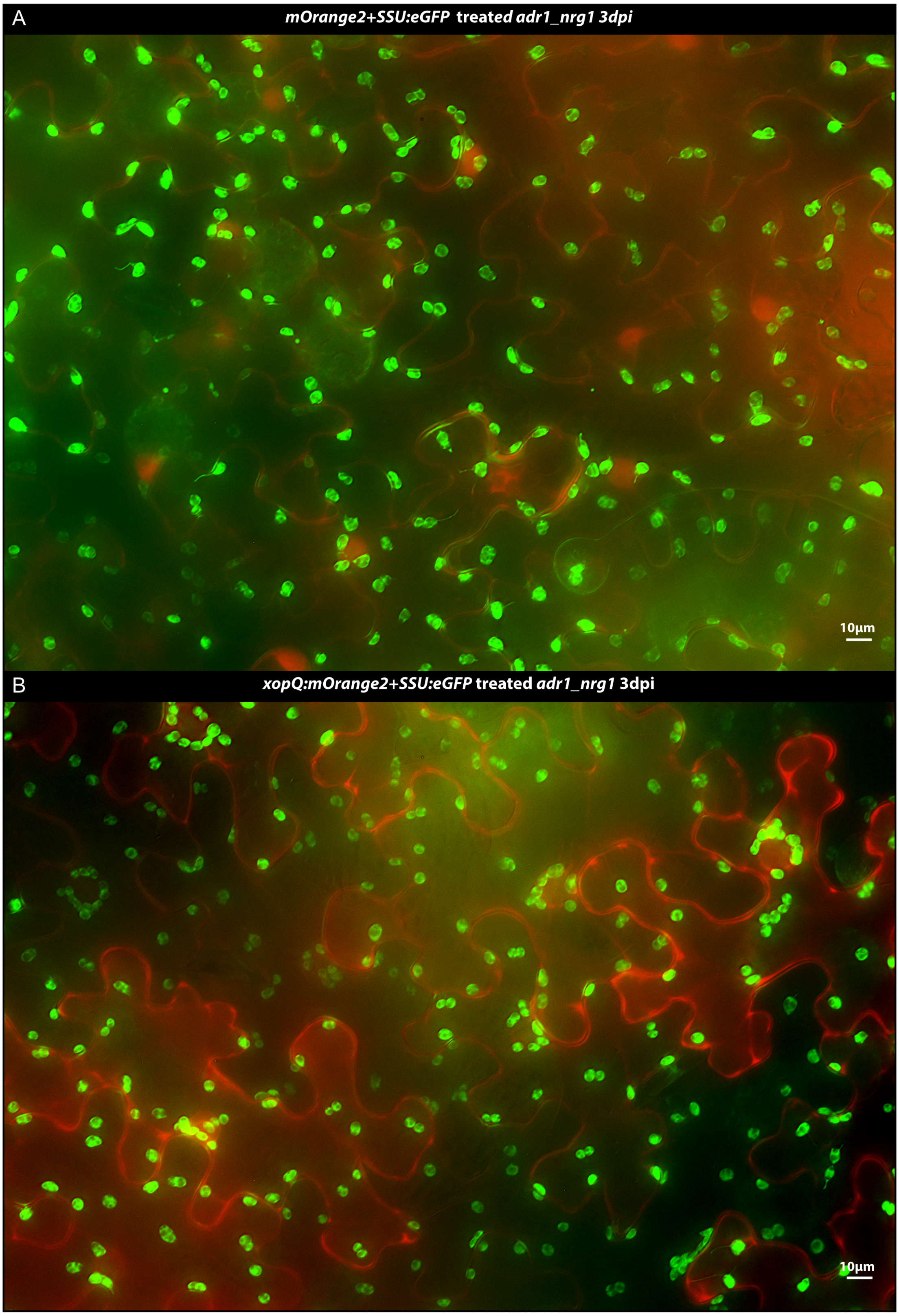
